# Isomerized Aβ in the brain can distinguish the status of amyloidosis in the Alzheimer’s Disease spectrum

**DOI:** 10.1101/2025.04.29.650793

**Authors:** Soumya Mukherjee, Reid Coyle, Celine Dubois, Keyla Perez, Catriona McLean, Colin L. Masters, Blaine R. Roberts

## Abstract

Extracellular amyloid plaques, the pathognomonic hallmark of Alzheimers Disease (AD), are also observed in cognitively unimpaired subjects in the preclinical stages. Progressive accumulation of fibrillar amyloid-β (Aβ) as plaques and perivascular deposits occur two decades prior to clinical onset, making Aβ a long-lived peptide. To characterize the amyloid plaques biochemically, both the Aβ-load as well the post-translational modifications (PTMs) could serve as markers for distinguishing the pre-clinical stage compared to later prodromal and clinical stages of AD. Recently, we described the presence of extensive isomerization of the Aβ N-terminus in AD post-mortem brains that are significantly increased compared to the age-matched non-AD control brains with Aβ aggregates in the frontal cortex. In this report, we used targeted mass spectrometry to conduct a quantitative analysis of the most common PTMs associated with Aβ; pyroglutamation, citrullination, N-terminal truncation (Aβ_4-x_), C-terminal truncation (Aβ_42_ and Aβ_40_), and isomerization of aspartic acid residues (Asp-1 and Asp-7) in postmortem human brain tissue from pathologically negative (no Aβ plaques) controls, controls with Aβ plaques, Parkinson’s disease (PD) with and without Aβ accumulation/plaques and symptomatic AD. The AD cases contained statistically significant amounts of Asp-1and Asp-7 isomerized Aβ_1-15_ (∼ 90 %) compared to controls (preclinical AD) and PD brains with fibrillar Aβ aggregates/deposits. We find that ratio of isomerized N-terminus Aβ (Aβ_1-15_) species in the brain detergent soluble pool differentiates older fibrillar Aβ deposits in symptomatic AD brain compared to Aβ deposits detected in preclinical AD and PD. Citrullinated Aβ_3pglu-15_ was increased only in symptomatic AD, highlighting this Aβ PTM is a unique feature of parenchymal plaques in advanced AD. Our results have implications for early therapeutic targeting of these modified species as well potential for better biofluid biomarker development for drug efficacy monitoring.

## Introduction

Alzheimer disease (AD) is most common form of dementia affecting more than 60 million people globally. Two pathological hallmarks of AD are extracellular amyloid-β (Aβ) deposits and intraneuronal fibrillary tangles composed of hyperphosphorylated tau.^1–3^ In the amyloidogenic pathway, sequential cleavages of the amyloid precursor protein (APP) by the β-secretase and γ-secretase lead to generation of Aβ peptides that are 37-43 amino acids long.^4^ Other than sporadic late-onset AD, multiple autosomal dominant mutations on the *APP* (Amyloid-β precursor protein) or *PSEN1* and *PSEN2* (Presenilin-1 and 2) genes are known to drive early-onset of the disease pathophysiology in autosomal dominantly inherited AD (ADAD).^5,6^ PET imaging as well as the fluid biomarkers (cerebrospinal fluid and blood) have established that amyloid deposition starts almost two-decades before symptom onset and cognitive deterioration in both sporadic late-onset AD (LOAD) as well as early-onset ADAD.^7–11^ Deposition of Aβ peptides as fibrillar aggregates in the brain correlates with its decreased clearance in AD, thereby making Aβ, a long-lived peptide.^12–14^

Multiple quantitative mass spectrometric studies of the parenchymal amyloid plaques have demonstrated a heterogeneous Aβ proteoform distribution, with Aβ_42_ being the most abundant species.^15–17^ Plaque associated Aβ peptides from AD brains are well known to undergo multiple N-terminal truncations, phosphorylation,^18^ oxidation, pyroglutamate formation,^19,20^ citrullination of the arginine residue,^21^ nitration,^22^ dityrosine cross-linking^23^ along with the significant amount of non-enzymatic isomerization at Asp-1 and Asp-7.^24–27^ The isomerization of Asp residues at the N-terminus (i.e. Asp-1 and Asp-7) has been described as the most common and extensive modification associated with Aβ aggregates in the AD brain.^17,24^ Using ultra high-performance liquid chromatography (UHPLC) coupled with drift tube ion mobility mass spectrometry (DTIM-MS), we have demonstrated Asp residues in the N-terminus of Aβ (Aβ_1-15_) extracted from the insoluble extracellular plaques was ∼ 85 % isomerized, compared to ∼ 50 % found in the age matched controls with amyloid plaque pathology.^17,28^ Abundant amyloid deposits of Aβ are observed in preclinical AD in the brains of individuals without any cognitive impairment, especially as a function of age.^29^ Recent cryo-EM (cryo-electron microscopy) imaging of amyloid plaque derived Aβ filaments demonstrate that the first eight to eleven residues are disordered at the N-terminus, while the protofilament core includes the residues from Tyr10 to Ala42.^30^ These imaging results support spontaneous chemical modification of the N-terminus Asp residues leading to the extensive isomerization. Recent biochemical and histopathological study on a postmortem tissue from a diverse cohort also highlighted the significance of isoAsp7 modification as a highly abundant Aβ PTM across different clinical conditions with Aβ aggregates including dementia with Lewy bodies, vascular dementia and symptomatic AD.^27^ With the recent success in anti-Aβ immunotherapy,^31,32^ that target the disordered N-terminus region of Aβ, indicates that a detailed understanding of PTMs in this region should yield better diagnostics and therapeutics for AD.

In this current study, using a well preserved frozen human brain tissues and with detailed pathological characterization, we extracted the detergent soluble fibrillar Aβ aggregates to address the question what Aβ PTMs are associated with AD neuropathologic changes (ADNC) in symptomatic AD, preclinical AD and non-AD dementia. Using a targeted mass spectrometric proteomic method we evaluated multiple isoforms of Aβ (both C-terminus as well as N-terminus truncations) from post-mortem brain tissue from advanced AD and compared them to age-matched non-demented individuals without Aβ plaques pathology (CU-AN) and pathologically positive controls with Aβ plaque deposits (CU-AP). We also investigated brains with Lewy body dementia and clinically diagnosed Parkinson’s disease (PD) with co-incidence of Aβ plaques (PD-AP) and PD brains without Aβ plaque pathology (PD-AN). Along with Asp-1 and Asp-7 isomerization we also measured the multiple PTMs (pyroglutamate and citrullination) associated with fibrillar Aβ aggregates at the N-terminus of Aβ. We first investigated if the Aβ isoforms and their absolute quantity could differentiate AD compared to CU-AP and PD-AP and correlated the biochemical measures with neuropathological scores. We also compared the N-terminus isomerized species of fibrillar Aβ and their ratios in CU-AP and PD-AP with that found in late symptomatic stage of AD.

## Materials and Methods

All LC-MS grade solvents, acetonitrile (ACN), formic acid (FA), trifluoroacetic acid (TFA), acetic acid, isopropanol, and urea, 3-[(3-cholamidopropyl)dimethylammonio]-1-propanesulfonate (CHAPS), sodium deoxycholate, iodoacetamide (IAA), tri-ethyl ammonium bicarbonate (TEAB) buffer, protease inhibitor cocktail (Complete EDTA free), NaCl, Tris buffers were purchased from Merck-Sigma or ThermoFischer Scientific. MS grade metalloprotease LysN from Grifola frondosa and Bond-Breaker™ TCEP Solution, neutral pH, were purchased from ThermoFischer Scientific. Biomasher was purchased from Omni International. The MS vials, Advanced Bio Peptide Mapping C18 Column (2.16 mm x 150 mm, 2.7 µm) and ESI low concentration tune mix used for instrument calibration were obtained from Agilent Technologies (Santa Clara, USA). Oasis HLB µElution 96 well-plates were purchased from Waters. Full-length protein ^15^N labelled tau1-441 (2N4R) was purchased from rPeptide (Georgia, USA). Amino acid analysed Lys-N peptides Aβ peptides DAEF(R+10)HDSGYEVHHQ, F(R+10)HDSGYEVHHQ, (K+8)GAIIGLMVGGVV, (K+8)GAIIGLMVGGVVIA, K(+8)LVFFAEDVGSN were purchased from New England Peptides. Stock solutions of SIS Aβ peptides were prepared in 2 % ACN, 0.05 % TFA to a final concentration of 200 fmol/µL and stored at -80 °C. All the isomerized and citrullinated Aβ peptide standards were commercially synthesized and purchased from JPT Peptide Technologies (Germany). All the SIS Aβ peptides were resuspended in 30 % ACN, 0.1 % FA at 0.2 nmol/µL which were subsequently diluted to ∼ 2 pmol/µL in 15 % ACN, 0.1 % FA and stored at -80 °C.

### Human brain tissues

Postmortem brain tissue samples were obtained from the Victorian Brain Bank (Australia). The cohort consisted of age-matched AD brains (n=59), Parkinson brains (n = 57) and age matched control brains (61). The brains were histopathologically analysed to determine the number of plaques and tangles. The AD brains met the standard criteria for AD neuropathological diagnosis (Table S1).

### Detergent soluble protein extraction and protein digestion

Approximately 50-80 mg of grey matter from postmortem frontal cortex tissues were homogenized in 200 µL of urea-detergent lysis buffer (8M Urea, 1 % CHAPS, 0.2 % sodium deoxycholate, 100 mM TEAB, pH 8.5, EDTA free protease inhibitor) using a probe sonicator (Branson, Sonifier® Cell Disruptor) in a series of 5 seconds at 30% power until complete homogenization (Figure 1A). The starting brain homogenate was centrifuged (Eppendorf microcentrifuge, 5415D) at 16,000 *g* for 20 mins at 4°C. The supernatant was transferred to fresh tubes and protein concentration was estimated using BCA assay. Protein content in each sample was determined using the BCA protein assay, and 25 µg of total protein was used for Lys-N digestion (typically 5 µL).

**Figure 1.**
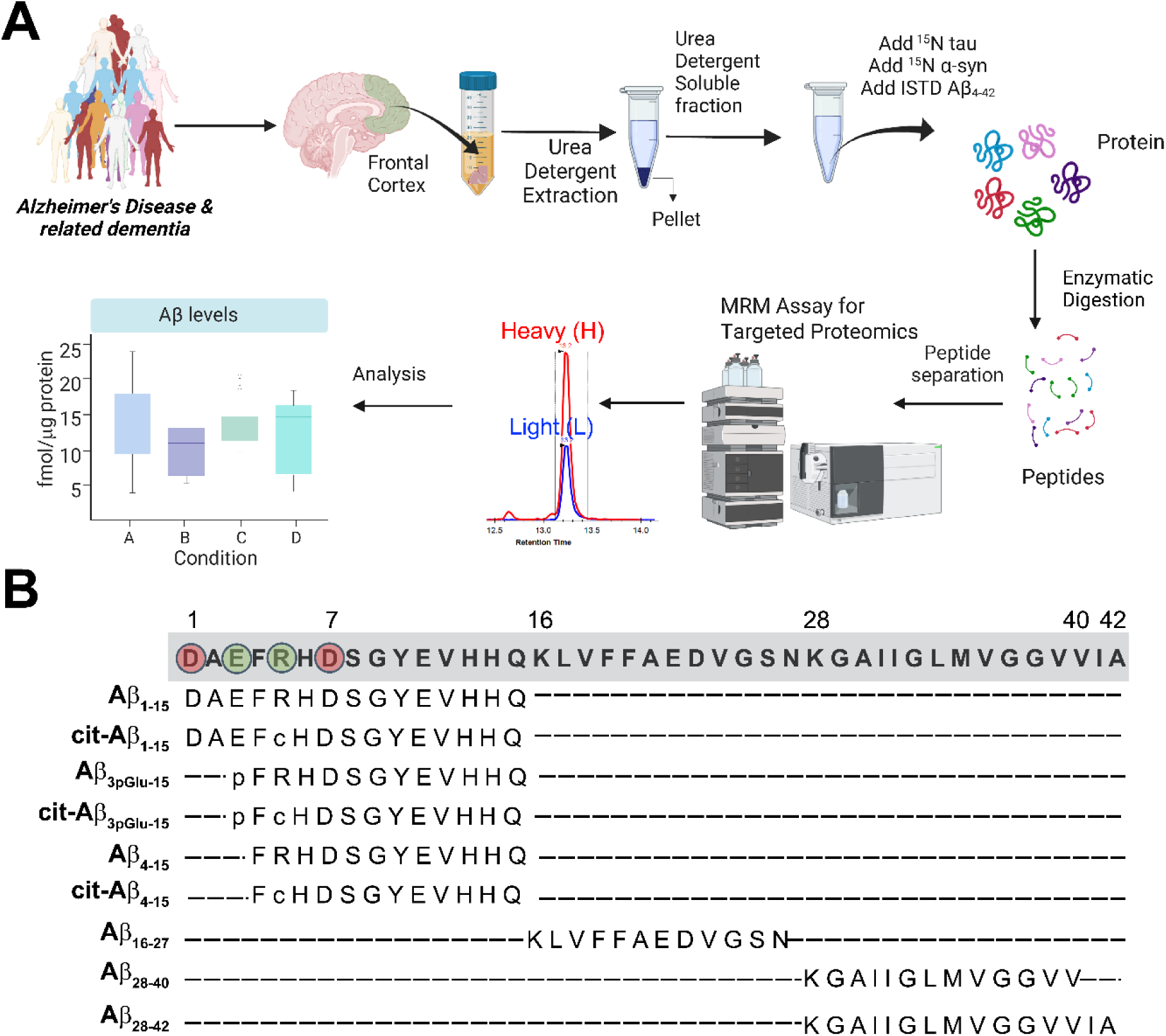
**(A)** Schematic workflow depicting the workflow for quantitative biochemical analysis of the levels of Aβ variants, tau isoforms and α-synuclein found in the frozen brain tissues of patients diagnosed with symptomatic AD, preclinical AD, Parkinson’s disease and cognitively unimpaired controls using targeted mass spectrometry. Created with Biorender.com (B) Schematic diagram of the Aβ peptide and the specific variants (Lys-N digested peptides) that were investigated in this study in the brain cohort using targeted proteomic assay. Aβ PTMs investigated: p, pyroglutamate; c, citrullination. Aspartic acid residues Asp-1 and Asp-7 are highlighted in red circle were further investigated for their isomerization ratio in the brains.

To estimate the absolute quantity of Aβ, tau and α-synuclein, a modified LysN-digestion workflow was used.^17,21^ Briefly, predetermined amounts 17 pmoles of isotopically labelled Aβ_4-42_ (R (^13^C6,^15^N4) and K (^13^C6, ^15^N2)), 10 ng of full-length ^15^N labelled tau (2N4R) and 1.25 ng ^15^N labelled full-length α-synuclein standard were add to 25 µg of total protein of the brain lysate. The brain lysates were then diluted with 25 µL 8M urea, 5 mM TCEP, 50 mM TEAB pH 8.5 buffer and sonicated for 5 minutes followed by incubation at 54 °C for 30 minutes. After reduction, the samples were cooled and alkylated with 20 mM final concentration iodoacetamide (IAA) in the dark for 30 minutes at 37 °C. The samples were diluted to 200 µL with 50 mM TEAB pH 8.5, 1.1 mM CaCl_2_ buffer to reduce the urea concentration to 1.2 M. The protein samples were digested with LysN enzyme (1:75 enzyme: protein ratio) overnight at 37 °C. The digested peptides were quenched with 10 % TFA to 0.1 % TFA in the solution and these acidified samples, stable isotope labelled LysN specific Aβ peptides were added as previously described.^17,21,33^ These samples were desalted using Oasis µElution HLB plates (Waters) and the eluent was lyophilized and stored at -80 °C prior mass spectrometry analysis.

To estimate phosphorylated tau species an aliquot of the brain lysate was subjected to trypsin digestion as previously described.^33^ Briefly, 25 µg of total brain lysates were denatured, reduced and alkylated as above. Then the samples were diluted with 200 µL with 50 mM TEAB pH 8.5, 1.1 mM CaCl_2_ buffer to reduce the urea concentration < 1 M and digested with 50 ng/µL trypsin enzyme (1:100 enzyme: protein ratio) overnight at 37 °C. Following digestion stable isotope labelled tau peptides pT181 (TPPAKT(+79.99)PPSSGEPPK(^13^C6, ^15^N2)), T181(TPPSSGEPPK(^13^C6, ^15^N2)), pT217 (TPSLPT(+79.99)PPTR(^13^C6, ^15^N4)) and T217 (TPSLPTPPTR(^13^C6, ^15^N4)) were added for phospho tau quantitation.^33^ The digested peptides were quenched, desalted, lyophilized as above and stored at -80 °C prior mass spectrometry analysis.

### Multiple reaction monitoring (MRM) of proteolytic peptides and data analysis

An Agilent 1200 Infinity series UHPLC system connected to 6495 QQQ (Agilent Technologies, USA) was used for the LC-ESI-QQQ-SRM assay (Figure 1A). Mobile phase A consisted of 0.1 % FA in water and mobile phase B of 0.1 % FA in 100 % ACN. Samples were run in randomized order to minimize bias. For absolute Aβ, tau and α-synuclein quantitation, the lyophilized Lys-N digested samples were resuspended in 50 µL 2 % ACN, 0.05 % FA buffer and 10 µg of Lys-N digested peptides were separated using an Agilent Advanced Bio Peptide Mapping C18 Column (2.1 mm x150 mm, 2.7 µm) maintained at 55°C in column compartment and eluted at 0.4 mL/min flowrate with the following gradient, 2.5 % B, 0 min; 6 % B, 5 min; 9 % B, 20 min; 22% B, 25 min; 29 % B, 35 min; 34 % B, 37 min; 81 % B, 38 min; 81 % B, 40 min; 2.5 % B, 41 min with a post-run equilibration for 2 min. The source ESI parameters as well as the collision energies were optimized for these peptides in the positive ion mode in Skyline software (20.2.1.315, MacCoss Lab, Department of Genome Sciences, University of Washington, Seattle, WA) and the method was imported into Agilent Mass Hunter Workstation (version 10.0.142) for data acquisition. The instrument parameters were the following: gas temperature 200°C, gas flow 15 L/min, nebulizer 40 psi, sheath gas temperature 250°C, and sheath gas flow 11 L/min. The capillary voltage was 4500 V and the nozzle voltage was set at 1000 V. The optimized iFunnel parameters were 200 V and 110 V for high and low-pressure RF respectively. The targeted QQQ data were imported into Skyline with formula annotations of the targeted peptides added to the method. The peak area for the Lys-N peptides from heavy labelled tau and α-synuclein were used in this study were compared to their heavy analogues (^15^N metabolically labelled total tau and α-synuclein), while Aβ_1-15_, Aβ_3pGlu-15_, cit-Aβ_3pGlu-15_, Aβ_4-15_, Aβ_16-27_, Aβ_28-40_ and Aβ_28-42_ peptides were compared to their heavy analogues (R=^13^C_6_, ^15^N_4_, K=^13^C_6_,^15^N_2_) to derive the light-to-heavy ratios. The absolute quantification was determined by using the known concentrations of the spiked in of the heavy labelled peptides. Individual isomerized Aβ_1-15_ was quantified by using the known concentrations of the spiked respective isomerized Aβ_1-15_ SIS peptides.^17^ Percentage isomerization on respective Aβ isomers was calculated as the fraction of isomerized peptide divided by the sum of unmodified and isomerized peptide pairs.

### Multiple reaction monitoring (MRM) of phospho-tau quantification

The same LC-MS system as above was used with a modified gradient; 2.5 % B, 0 min; 25 % B, 15 min; 34 % B, 17 min; 81 % B, 17.2 min; 81 % B, 18.75min; 2.5 % B, 18.8 min. Data for non-phosphorylated and phosphorylated tau peptides were extracted in the Skyline software using the transition lists and co-elution of the heavy stable isotope labeled (SIS) spiked standard. The peak area for the tryptic tau/phospho-tau peptides were compared to their heavy analogues (labelled total tau R=^13^C_6_, ^15^N_4_, K=^13^C_6_,^15^N_2_ for phospho-tau SIS peptides) and the absolute quantification was determined by using the known concentrations of the spiked in SIS peptides.

### Statistical analysis

Data analyses were performed with R and GraphPad Prism. To determine statistically significant differences between potential biomarkers, the adjusted *P* values were calculated using unpaired non-parametric one-way analysis of variance (ANOVA), corrected for multiple comparison false discovery rate (*P* < 0.05) with Benjamini–Hochberg correction.^34^ A value of *p*<□0.05 was considered as statistically significant. Data are represented as mean ± standard deviation (SD). The diagnostic performance of the biomarker was assessed by the receiver operating curves (ROC) and area under the curve (AUC). The diagnostic accuracy of each biomarker was described by the ROC curve and summarized by the AUC with 95 % confidence interval (CI). Correlations of the biomarkers with ADNC scores and age were calculated using Pearson correlations.

### Ethics approval

Fresh frozen brain samples were obtained from the Victorian Brain Bank (VBB). The study was approved by the ethics committee of the University of Melbourne (Ethics 1750801.3).

## Results

### Total Aβ and modified Aβ species in human brains

Using targeted mass spectrometry, we quantified Aβ peptides from the detergent soluble brain homogenates.^17,33^ We measured the concentration of two canonical C-terminus Aβ species (Aβ_42_ and Aβ_40_), mid-domain (Aβ_16-27_) and sequentially N-truncated species (Aβ_1-15_, Aβ_4-15_, Aβ_3pGlu-15_) and associated post-translational modifications (PTMs) (Figure 1B) in a cohort of age-matched individuals with Alzheimer’s disease (AD, n = 60), non-demented controls with Aβ plaques (CU-AP, n = 35) and controls negative for Aβ plaque pathology (CU-AN, n = 23) and Parkinson’s disease (PD) patients with Aβ plaque co-pathology (PD-AP, n = 30) and PD patients negative for Aβ plaque pathology (PD-AN, n = 28) (Table 1). The parenchymal Aβ plaques are primarily composed of Aβ_42_ and is known to highly correlate with the mid-domain Aβ peptide, the most utilized marker for amyloid aggregates/fibrils in brain homogenates for proteomic analysis.^15,17,34^ Aβ_42_ concentration was 10-fold higher than Aβ_40_ across the whole cohort (Figure 2A-B, Table 2). We observed Aβ_42_ was significantly increased (*p* < 0.0001) in AD (232.6 ± 186.9 fmol/µg protein) compared to CU-AN (47.2 ± 36.7 fmol/µg protein), CU-AP (94.0 ± 145.9 fmol/µg protein), PD-AN (29.9 ± 25.6 fmol/µg protein) and PD-AP (158.6 ± 157.2 fmol/µg protein) (Table 2, Figure 2A). The Aβ_42_ levels correlated significantly (r = 0.94, *p* < 0.0001) with the concentration of Aβ mid-domain (Aβ_16-27_) across the whole detergent soluble brain cohort. We observed no significant difference of Aβ_42_ levels in the CU-AN (47.2 ± 36.7 fmol/µg protein) compared to CU-AP (94 ± 145.9 fmol/µg protein) (Figure 2A). No significant difference in the Aβ_42_ levels were observed between CU-AP and PD-AP (156.8 ± 157.2 fmol/µg protein). Aβ_40_ concentration was significantly increased in AD (16.4 ± 48.1 fmol/µg protein) compared to CU-AN (1.3 ± 1.9 fmol/µg protein) (*p* < 0.0001), CU-AP (0.8 ± 0.7 fmol/µg protein) (*p* < 0.0001), PD-AN (0.7 ± 0.9 fmol/µg protein) (P < 0.0001) (Table 2, Figure 2B). We observed significant differences in Aβ_16-27_ levels between CU-AP (92.5 ± 153.8 fmol/µg protein) and CU-AN (15.6 ± 28.8 fmol/µg protein), as well as between PD-AP (150.4 ± 163.4 fmol/µg protein) and PD-AN (9.6 ± 15.9 fmol/µg protein) (Figure 2C, Table 2). In addition, Aβ_40/42_ was significantly different between CU-AN (0.03 ± 0.06) and CU-AP (0.01 ± 0.01) (*p* = 0.013, Figure S2) and PD-AN (0.04 ± 0.05) and PD-AP (0.02 ± 0.04) (*p* = 0.0025, Figure S2). No significant difference for Aβ_40/42_ were observed between AD (0.06 ± 0.12) and CU-AN (Figure S2).

**Figure 2.**
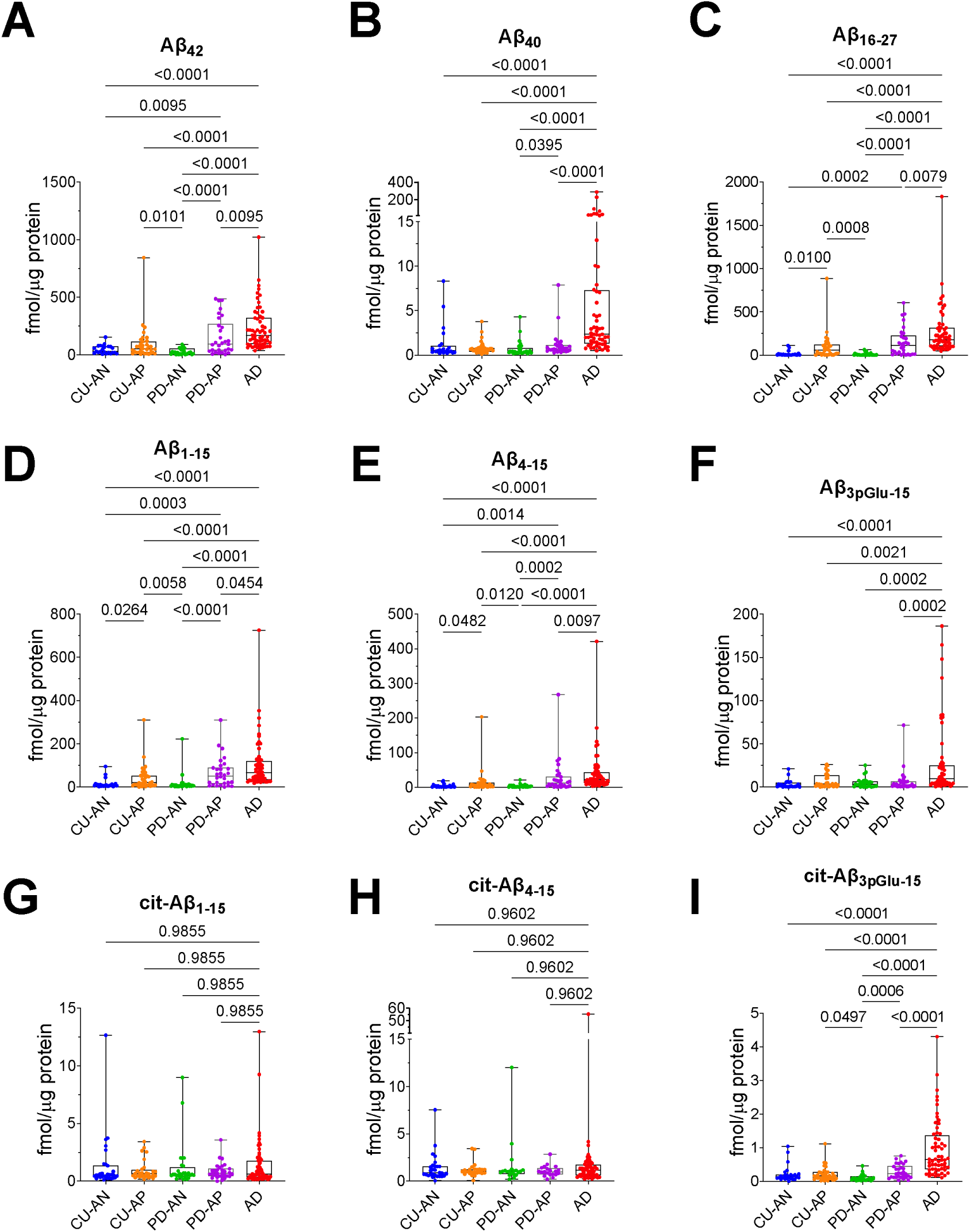
Box and whisker plots for the concentrations (fmol/µg protein) for Aβ species in the detergent soluble brain homogenates. (A) Aβ_28-42_ (B) Aβ_28-40_ (C) Aβ_16-27_ (D) Aβ_1-15_ (E) Aβ_4-15_ (F) Aβ_3pGlu-15_ (G) cit-Aβ_1-15_ (H) cit-Aβ_4-15_ and (I) cit-Aβ_3pGlu-15_. AD, Alzheimers disease (red); CU-AN, control brains without Aβ plaques (blue); CU-AP, control brains with Aβ plaques (orange); PD-AN, Parkinson’s disease (PD) brains without Aβ plaques (green); PD-AP, PD brains with Aβ plaques (purple). Statistically significant differences between the different groups are highlighted, Kruskal-Wallis test followed by Benjamini-Hochberg correction was performed for multiple comparisons.

**Table 1.** Demographics of the postmortem frontal cortex brain cohort including their Alzheimer’s disease neuropathologic changes (ABC score)^81^.

|  | AD | CU-AN | CU-AP | PD-AN | PD-AP |
| --- | --- | --- | --- | --- | --- |
| N | 60 | 23 | 35 | 28 | 30 |
| Sex (M/F) | (39/21) | (18/5) | (19/16) | (18/10) | (19/11) |
| Age (years) | 77.35 (7.11) | 73.72 (11.25) | 79.61 (7.69) | 76.78 (7.44) | 77.86 (6.66) |
| PMI (hours) | 37.74 (19.93) | 39.68 (12.74) | 39.57 (19.17) | 35.13 (15.85) | 36.12 (18.00) |
| A-score: n per stage<br>0/1/2/3 | 0/0/11/49 | 22/1/0/0 | 7/27/1/0 | 28/0/0/0 | 0/20/9/1 |
| B-score: n per stage<br>0/1/2/3 | 0/0/1/59 | 17/6/0/0 | 17/17/1/0 | 21/7/0/0 | 17/11/2/0 |
| C-score: n per stage<br>0/1/2/3 | 0/0/11/49 | 23/0/0/0 | 33/2/0/0 | 28/0/0/0 | 0.07 (0.25) |

All data are represented as mean values ( ± standard deviation (SD)); M, male; F, female; PMI, postmortem interval; A-score, Aβ /amyloid plaque score (Thal phases);<sup>82</sup> B-score, NFT score (Braak stage)<sup>83,84</sup>; C-score: Neuritic plaque score (CERAD)<sup>85</sup>.

**Table 2.**
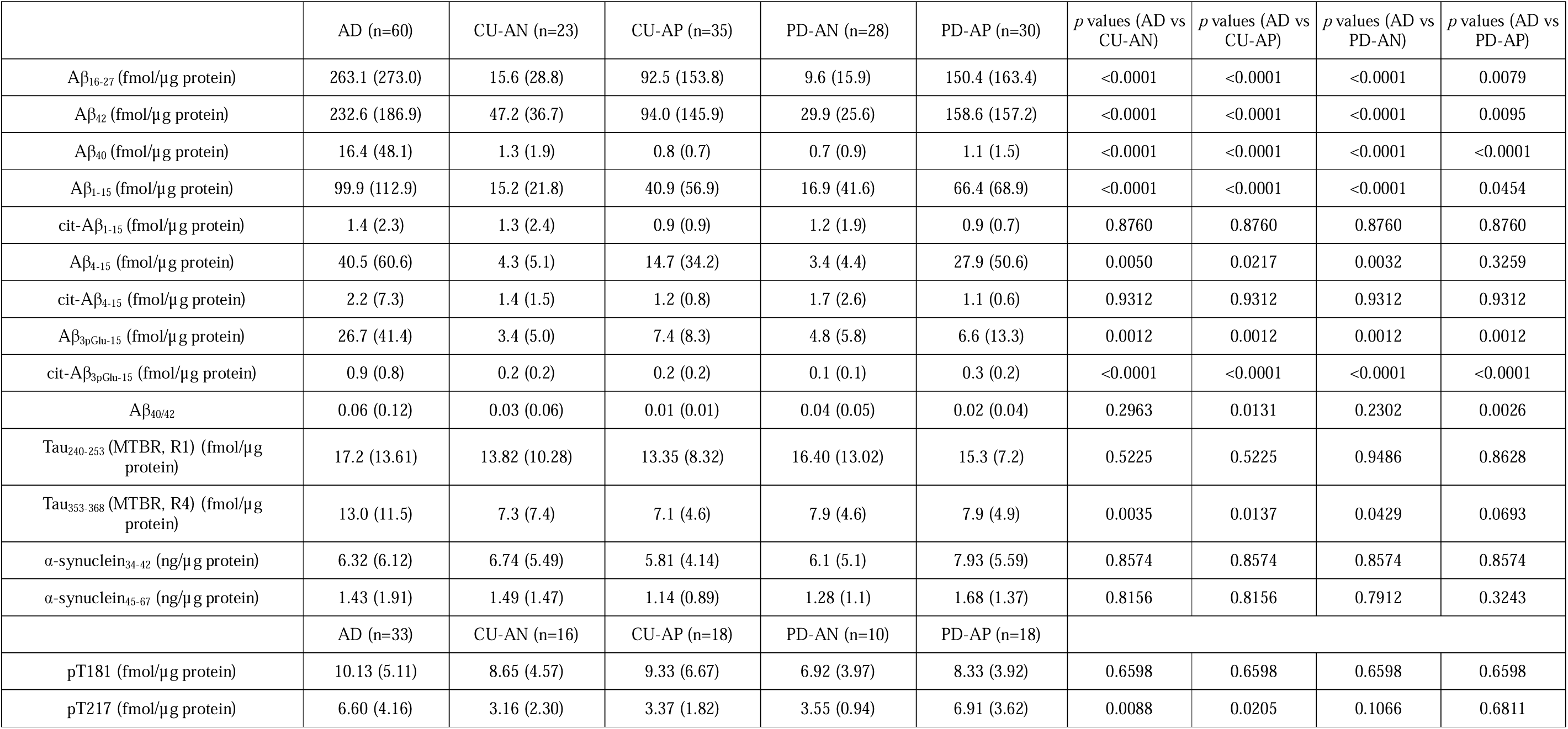
Quantitative estimates of detergent soluble Aβ isoforms, tau isoforms, phospho-tau and α-synuclein from the frontal cortex. All data are represented as mean values (± standard deviation (SD)); *p-*values are derived from Kruskal-Wallis test, corrected for multiple comparison false discovery rate with Benjamini-Hochberg correction.

|  | AD (n=60) | CU-AN (n=23) | CU-AP (n=35) | PD-AN (n=28) | PD-AP (n=30) | *p* values (AD vs CU-AN) | *p* values (AD vs CU-AP) | *p* values (AD vs PD-AN) | *p* values (AD vs PD-AP) |
| --- | --- | --- | --- | --- | --- | --- | --- | --- | --- |
| Aβ<sub>16-27</sub> (fmol/μg protein) | 263.1 (273.0) | 15.6 (28.8) | 92.5 (153.8) | 9.6 (15.9) | 150.4 (163.4) | <0.0001 | <0.0001 | <0.0001 | 0.0079 |
| Aβ<sub>42</sub> (fmol/μg protein) | 232.6 (186.9) | 47.2 (36.7) | 94.0 (145.9) | 29.9 (25.6) | 158.6 (157.2) | <0.0001 | <0.0001 | <0.0001 | 0.0095 |
| Aβ<sub>40</sub> (fmol/μg protein) | 16.4 (48.1) | 1.3 (1.9) | 0.8 (0.7) | 0.7 (0.9) | 1.1 (1.5) | <0.0001 | <0.0001 | <0.0001 | <0.0001 |
| Aβ<sub>1-15</sub> (fmol/μg protein) | 99.9 (112.9) | 15.2 (21.8) | 40.9 (56.9) | 16.9 (41.6) | 66.4 (68.9) | <0.0001 | <0.0001 | <0.0001 | 0.0454 |
| cit-Aβ<sub>1-15</sub> (fmol/μg protein) | 1.4 (2.3) | 1.3 (2.4) | 0.9 (0.9) | 1.2 (1.9) | 0.9 (0.7) | 0.8760 | 0.8760 | 0.8760 | 0.8760 |
| Aβ<sub>4-15</sub> (fmol/μg protein) | 40.5 (60.6) | 4.3 (5.1) | 14.7 (34.2) | 3.4 (4.4) | 27.9 (50.6) | 0.0050 | 0.0217 | 0.0032 | 0.3259 |
| cit-Aβ<sub>4-15</sub> (fmol/μg protein) | 2.2 (7.3) | 1.4 (1.5) | 1.2 (0.8) | 1.7 (2.6) | 1.1 (0.6) | 0.9312 | 0.9312 | 0.9312 | 0.9312 |
| Aβ<sub>3pGlu-15</sub> (fmol/μg protein) | 26.7 (41.4) | 3.4 (5.0) | 7.4 (8.3) | 4.8 (5.8) | 6.6 (13.3) | 0.0012 | 0.0012 | 0.0012 | 0.0012 |
| cit-Aβ<sub>3pGlu-15</sub> (fmol/μg protein) | 0.9 (0.8) | 0.2 (0.2) | 0.2 (0.2) | 0.1 (0.1) | 0.3 (0.2) | <0.0001 | <0.0001 | <0.0001 | <0.0001 |
| Aβ<sub>40/42</sub> | 0.06 (0.12) | 0.03 (0.06) | 0.01 (0.01) | 0.04 (0.05) | 0.02 (0.04) | 0.2963 | 0.0131 | 0.2302 | 0.0026 |
| Tau<sub>240-253</sub> (MTBR, R1) (fmol/μg protein) | 17.2 (13.61) | 13.82 (10.28) | 13.35 (8.32) | 16.40 (13.02) | 15.3 (7.2) | 0.5225 | 0.5225 | 0.9486 | 0.8628 |
| Tau<sub>353-368</sub> (MTBR, R4) (fmol/μg protein) | 13.0 (11.5) | 7.3 (7.4) | 7.1 (4.6) | 7.9 (4.6) | 7.9 (4.9) | 0.0035 | 0.0137 | 0.0429 | 0.0693 |
| α-synuclein<sub>34-42</sub> (ng/μg protein) | 6.32 (6.12) | 6.74 (5.49) | 5.81 (4.14) | 6.1 (5.1) | 7.93 (5.59) | 0.8574 | 0.8574 | 0.8574 | 0.8574 |
| α-synuclein<sub>45-67</sub> (ng/μg protein) | 1.43 (1.91) | 1.49 (1.47) | 1.14 (0.89) | 1.28 (1.1) | 1.68 (1.37) | 0.8156 | 0.8156 | 0.7912 | 0.3243 |
|  | AD (n=33) | CU-AN (n=16) | CU-AP (n=18) | PD-AN (n=10) | PD-AP (n=18) |  |  |  |  |
| pT181 (fmol/μg protein) | 10.13 (5.11) | 8.65 (4.57) | 9.33 (6.67) | 6.92 (3.97) | 8.33 (3.92) | 0.6598 | 0.6598 | 0.6598 | 0.6598 |
| pT217 (fmol/μg protein) | 6.60 (4.16) | 3.16 (2.30) | 3.37 (1.82) | 3.55 (0.94) | 6.91 (3.62) | 0.0088 | 0.0205 | 0.1066 | 0.6811 |

The most abundant N-terminus Aβ peptide in the detergent soluble brain fractions was Aβ_1-15_, correlating positively with Aβ_16-27_ (r = 0.90, *p* < 0.0001) and Aβ_42_ (r = 0.83, *p* < 0.0001) across the whole cohort (Figure S1). Aβ_1-15_ was significantly increased in AD (99.9 ± 112.9 fmol/ µg protein) compared to CU-AN (15.2 ± 21.8 fmol/ µg protein) (*p* < 0.0001), CU-AP (40.9 ± 56.9 fmol/µg protein) (*p* < 0.0001), PD-AN (16.9 ± 41.6 fmol/ µg protein) (*p* < 0.0001), and PD-AP (66.4 ± 68.9fmol/µg protein) (*p* = 0.0454) (Table 2, Figure 2D). We further observed significant differences between PD-AN and PD-AP (*p* < 0.0001) (Figure 2D). The next abundant N-truncated amyloid species was the Aβ_4-15_ (Figure 2E) and accounted for a one-fourth of total amyloid aggregates/deposits extracted from these brains (compared to the levels of Aβ_16-27_). Aβ_4-15_ was significantly higher in AD (40 ± 60.6 fmol/ µg protein) compared to CU-AP (14.7 ± 34.2 fmol/µg protein) (*p* = 0.0217), CU-AN (4.3 ± 5.1 fmol/ µg protein) (*p* = 0.005) and PD-AN (3.4 ± 4.4 fmol/ µg protein) (*p* = 0.0032) (Figure 2E, Table 2). No significant change was observed for Aβ_4-15_ in PD-AP (27.9 ± 50.6 fmol/ µg protein) compared to AD (40.5 ± 60.6 fmol/ µg protein). We investigated N-terminus pyroglutamate modified peptide Aβ_3pGlu-15_, and observed this modified peptide was five times lower than Aβ_1-15_ (Table 2, Figure 2F). Aβ_3pGlu-15_ was significantly higher in AD brains (26.7 ± 41.4 fmol/µg protein) compared to PD-AP (6.6 ± 13.3 fmol/µg protein ) (*p* = 0.0012), CU-AP (7.4 ± 8.3 fmol/µg protein ) (*p* = 0.0012), PD-AN (4.8 ± (5.8) fmol/ µg protein ) (*p* = .0012), and CU-AN (1.4 ± 1.5 fmol/µg protein ) (*p* = 0.0012) (Table 2). Our results further indicated that while Aβ_1-15_ and Aβ _4-15_ are highly correlated (r = 0.87, *p* < 0.0001), Aβ_pGlu3-15_ poorly correlates with both Aβ_1-15_ (r = 0.40, *p* < 0.0001) and Aβ _4-15_ (r = 0.41, *p* < 0.0001) (Figure S1).

Recently, we described a novel PTM on Aβ, citrullination of Arg5 residue, derived from the extracellular Aβ fibrils/aggregates from AD brains.^21^ We noticed hypercitrullination of pyroglutamte modified Aβ in both sporadic and familial AD brains.^21^ In this study, we quantified the levels of citrullinated Aβ_1-15_, Aβ _4-15_, Aβ_3pGlu-15_ peptides in the detergent soluble brain homogenates (Figure 2G-I). We observed no significant changes in the levels of cit-Aβ_1-15_ and cit-Aβ_4-15_ in AD compared to other brain samples (CU-AN, CU-AP, PD-AN and PD-AP) (Figure 1G-H). We observed cit-Aβ_3pGlu-15_ was 2-orders of magnitude lower compared to total Aβ (Aβ_16-27_) (Figure 2I). We observed cit-Aβ_3pGlu-15_ was significantly increased in AD (0.9 ± 0.8 fmol/ µg protein, *p* < 0.0001) compared to CU-AN (0.2± 0.1 fmol/ µg protein), CU-AP (0.2 ± 0.2 fmol/ µg protein), PD-AN (0.1 ± 0.1 fmol/ µg protein) and PD-AP (0.3 ± 0.1 fmol/ µg protein) (Table 2, Figure 2I). Significant differences (*p* = 0.0006) were also observed for cit-Aβ_3pGlu-15_ between PD-AN and PD-AP (Figure 2I, Table 2). These results highlight that hypercitrullination of pyroglutamate modified Aβ is a feature of advanced Aβ aggregates in late symptomatic stages of AD.

In addition, we compared the AD neuropathologic changes with the biochemical Aβ isoforms measures and observed higher Aβ_42_, Aβ_40_, Aβ_16-27_, Aβ_1-15_ Aβ_4-15_ and Aβ_3pGlu-15_ levels in AD brains associated with higher Thal amyloid phases (A2 and A3) (Figure 3), while lower Aβ levels in CU-AN and PD-AN correlated with lower Thal amyloid phases (A0/A1). Aβ_42_, Aβ_40_, Aβ_16-27_, Aβ_1-15_ and Aβ_4-15_ levels in the CU-AP and PD-AP brains associated with the intermediate Thal amyloid phases (A1/A2) (Figure 3). Interestingly, CU-AP and PD-AP brains had either none or sparse (C0/C1) neuritic plaques compared to moderate to frequent neuritic plaques in AD brains (C2/C3) and this was reflected in positive correlations with Aβ_42_, Aβ_40_, Aβ_16-27_, Aβ_1-15_ Aβ_4-15_ and Aβ_3pGlu-15_ levels (Figure 3).

**Figure 3:**
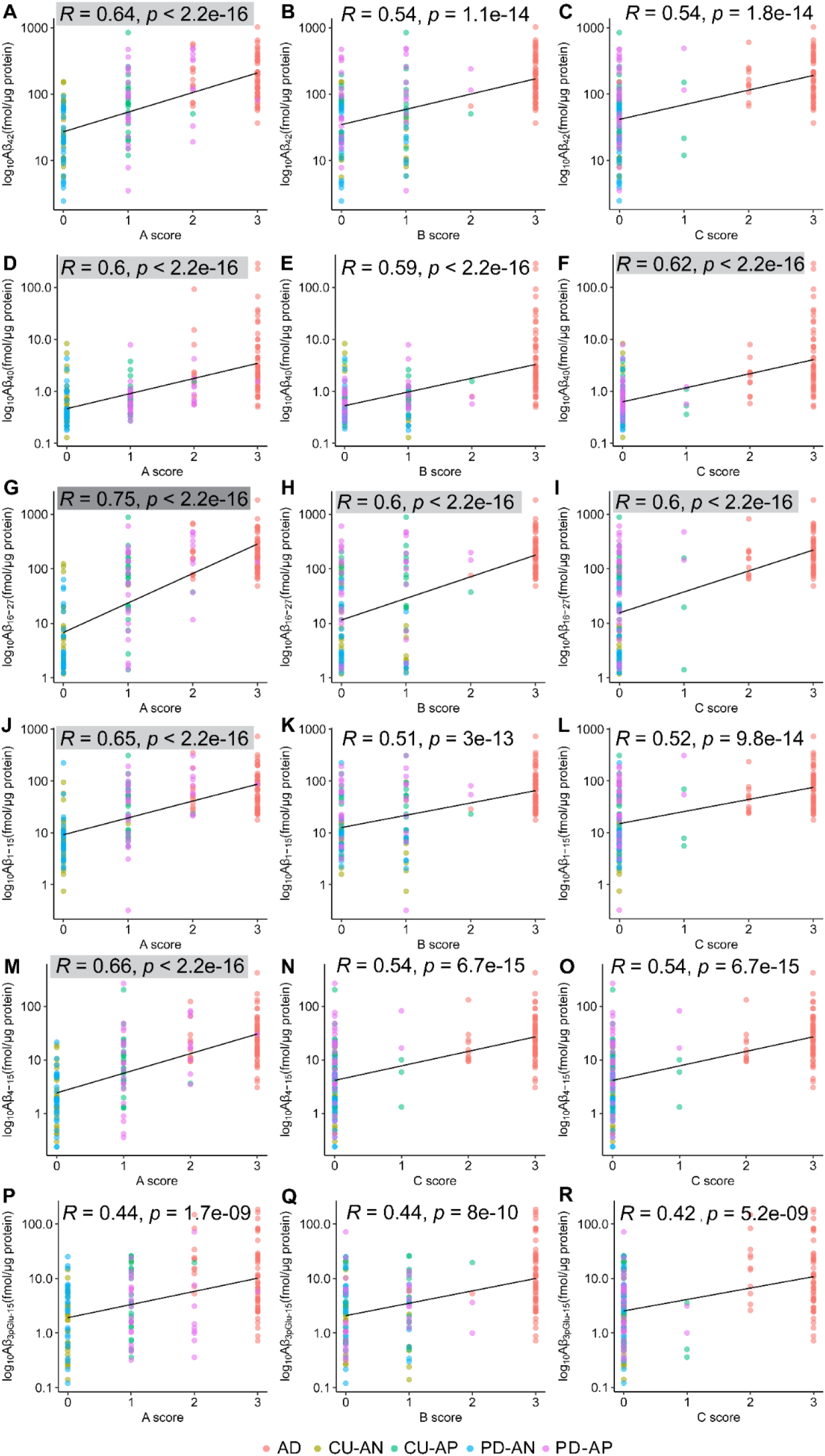
Pearson correlation analyses between the log_10_ transformed quantitative measures (fmol/µg protein) of Aβ_28-42_, Aβ_28-40_, Aβ_16-27_, Aβ_1-15_, Aβ_4-15_ and Aβ_3pGlu-15_ with neuropathological Thal amyloid A scores (A, D, G, J, M, N, P), Braak tau stages, B scores (B, E, H, K, N, Q) and neuritic plaques C scores (C, F, I, L, O, R). Strong correlation (highlighted in dark grey) was observed for Aβ_16-27_ (mid-domain) with Thal amyloid phases (A scores) and moderate correlations (highlighted in light grey) were observed with modified Braak tau scores and neuritic plaque scores. Moderate correlations (highlighted in light grey) were also observed for Aβ_28-42_, Aβ_28-40_, Aβ_1-15_ and Aβ_4-15_ with Thal amyloid phases.

### Distinguishing early and late ADNC stages using A**β** isoforms

Next, we investigated the overlap between advanced AD and early ADNC stages for the Aβ isoforms quantified in the detergent soluble brain fractions. We calculated the area under the curve (AUC) from the receiver operating characteristic (ROC) curves for the Aβ isoforms (Figure 4A-B). Aβ_42_ levels performed significantly better when distinguishing AD from brains without amyloid aggregates/fibrillar (CU-AN) deposits (AUC [95 % CI] =0.93 [0.87-0.98]) and PD-AN (AUC [95 % CI] = 0.98 [0.95-1.0]) (Figure 4A, Figure S4A). We observed Aβ_40_ accurately distinguished AD from for CU-AP (AUC = 0.90 [0.84-0.96]) and PD-AP (AUC = 0.86[0.77-0.94]) (Figure 4A, Figure S4B). Aβ_40_ also demonstrated slightly lower accuracy in distinguishing AD to CU-AN (AUC = 0.85 [0.75-0.95] compared to Aβ_42_ and Aβ_16-27_ (Figure 4A, Figure S4B). In addition, Aβ_16-27_ accurately distinguished AD from CU-AN (AUC = 0.98 [0.94-1.0]) and PD-AN (AUC = 0.99 [0.99-1]) (Figure 4A, Figure S4C). While Aβ_42_ and Aβ_16-27_ levels in the brain had lower performance when differentiating AD from CU-AP (AUC = 0.83[0.74-0.91] for Aβ_42_, AUC = 0.83[0.74-0.92] for Aβ_16-27_) and PD-AP (AUC = 0.66 [0.53-0.79] for Aβ_42_, AUC = 0.69 [0.56-0.81] for Aβ_16-27_) (Figure 4A, Figure S4A and Figure S4C).

**Figure 4.**
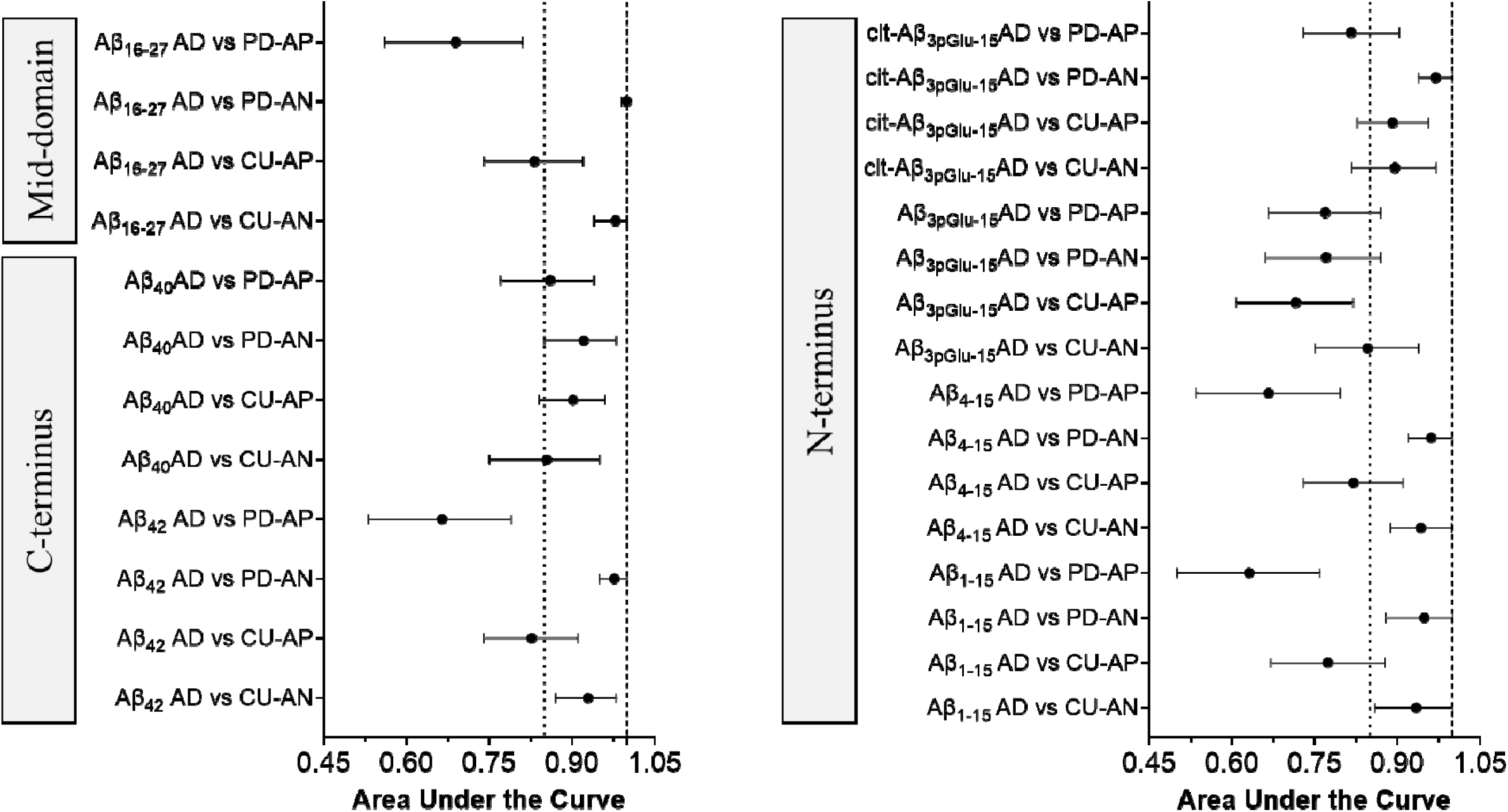
Correspondence with detergent soluble Aβ isoform levels in other cognitively unimpaired control brains, preclinical AD and co-morbidities compared to symptomatic AD. Correspondence of (A) C-terminus and mid-domain Aβ peptides and (B) N-terminus Aβ peptides in AD compared to other co-pathologies with and without brain Aβ aggregates/deposits. Points represent the receiver operating characteristic (ROC) from the area under the curve (AUC) estimate (middle point) and the error bars represent the 95 % confidence intervals. AD, Alzheimer’s disease; CU-AN, control brains without Aβ plaques; CU-AP, control brains with Aβ plaques; PD-AN, Parkinson’s disease (PD) brains without Aβ plaques; PD-AP, PD brains with Aβ plaques. Note Aβ_28-40_ as well as citrullinated Aβ_3pGlu-15_ have higher diagnostic performance in differentiating late ADNC compared to moderate ADNC.

We then assessed the diagnostic performance of N-terminus Aβ peptides. We observed Aβ_1-15_ and Aβ_4-15_ distinguished AD with high accuracy from CU-AN (AUC = 0.93 [0.86-1.0] for Aβ_1-15_, AUC = 0.94 [0.88-0.99] for Aβ_4-15_) and PD-AN (AUC = 0.95 [0.88-1.0] for Aβ_1-15_, AUC = 0.96 [0.92-1.0] for Aβ_4-15_) (Figure 4B, Figure S4D-E). In contrast, Aβ_1-15_ and Aβ_4-15_ had comparatively lower performance in distinguishing AD from CU-AP (AUC = 0.77 [0.67-0.88] for Aβ_1-15_, AUC = 0.82 [0.73-0.91] for Aβ_4-15_) (Figure 4B, Figure S4D-E). We observed slightly lower performance for Aβ_1-15_ and Aβ_4-15_ when discriminating AD from PD-AP (AUC = 0.63 [0.50-0.76] for Aβ_1-15_, AUC = 0.67 [0.54-0.80] for Aβ_4-15_) (Figure 4B, Figure S4D-E). In addition, Aβ_3pGlu-15_ showed lower accuracy in discriminating AD from CU-AP (AUC = 0.71 [0.61-0.82]) and PD-AP (AUC = 0.77 [0.66-0.87]) (Figure 4B, Figure S4F). More importantly, cit-Aβ_pGlu3-15_ showed very high accuracy in discriminating AD from CU-AN (AUC = 0.89 [0.82-0.97]) and PD-AN (AUC = 0.97 [0.94-1.0]) (Figure 4B, Figure S4G). In addition, cit-Aβ_pGlu3-15_ also accurately discriminated AD from CU-AP (AUC = 0.89 [0.83-0.96]) and PD-AP (AUC = 0.81[0.73-0.90) (Figure 4B, Figure S4G). The better diagnostic performance of cit-Aβ_pGlu3-15_ in separating early/intermediate stages of the AD compared to late ADNC indicates hypercitrullination is more closely associated with mature/older plaques found in symptomatic AD brains.

### Quantitation of tau, p-tau and **α**-synuclein in the detergent soluble fraction

Phosphorylated tau and certain species of from the microtubule binding region (MTBR) are known to be increased in AD brains compared to preclinical AD.^33,35,36^ We measured the total tau levels using proline rich region (PRR) and MTBR species along with phosphorylated tau peptide (p-tau 181 and p-tau 217) levels in the detergent soluble brain homogenates using targeted mass spectrometry (Figure 5 and Figure S5). MTBR tau species (353-368 residue, R4) was significantly increased in AD (13.00 ± 11.50 fmol/µg protein) compared to CU-AN (7.30 ± 7.40 fmol/µg protein) (*p* = 0.0035), CU-AP (7.10 ± 4.58 ng/µg protein) (*p* = 0.0137) and PD-AN (7.90 ± 4.88 ng/µg protein) (*p* = 0.0429) (Figure 5A). We found no significant changes for the phospho tau 181 (pT181) between AD and other comorbidities (Figure 4B). Phosphorylated tau 217 (pT217) was significantly increased in AD (6.6 ± 4.1 fmol/µg protein) compared to CU-AN (3.1 ± 2.3 fmol/µg protein) (*p* = 0.0088), CU-AP (3.4 ± 1.8 fmol/µg protein) (*p* = 0.0088) and PD-AN (3.5 ± 0.9 fmol/µg protein) (*p* = 0.0205) (Figure 5C). We also assessed the diagnostic performance of pT217 in detergent soluble fraction (Figure S6). We observed pT217 distinguished AD with modest accuracy from CU-AN (AUC = 0.76 [0.62-0.90]), PD-AN (AUC = 0.69 [0.54-0.85]) and CU-AP(AUC = 0.72 [0.60-0.87]) (Figure S6).We found pT217 levels did not differentiate between AD and PD-AP (AUC = 0.54 [0.34-0.71]) (Figure S6). We also measured detergent soluble α-synuclein levels in this cohort, however there were no statistical changes in the levels across the different groups (Table 2 and Figure S7).

**Figure 5.**
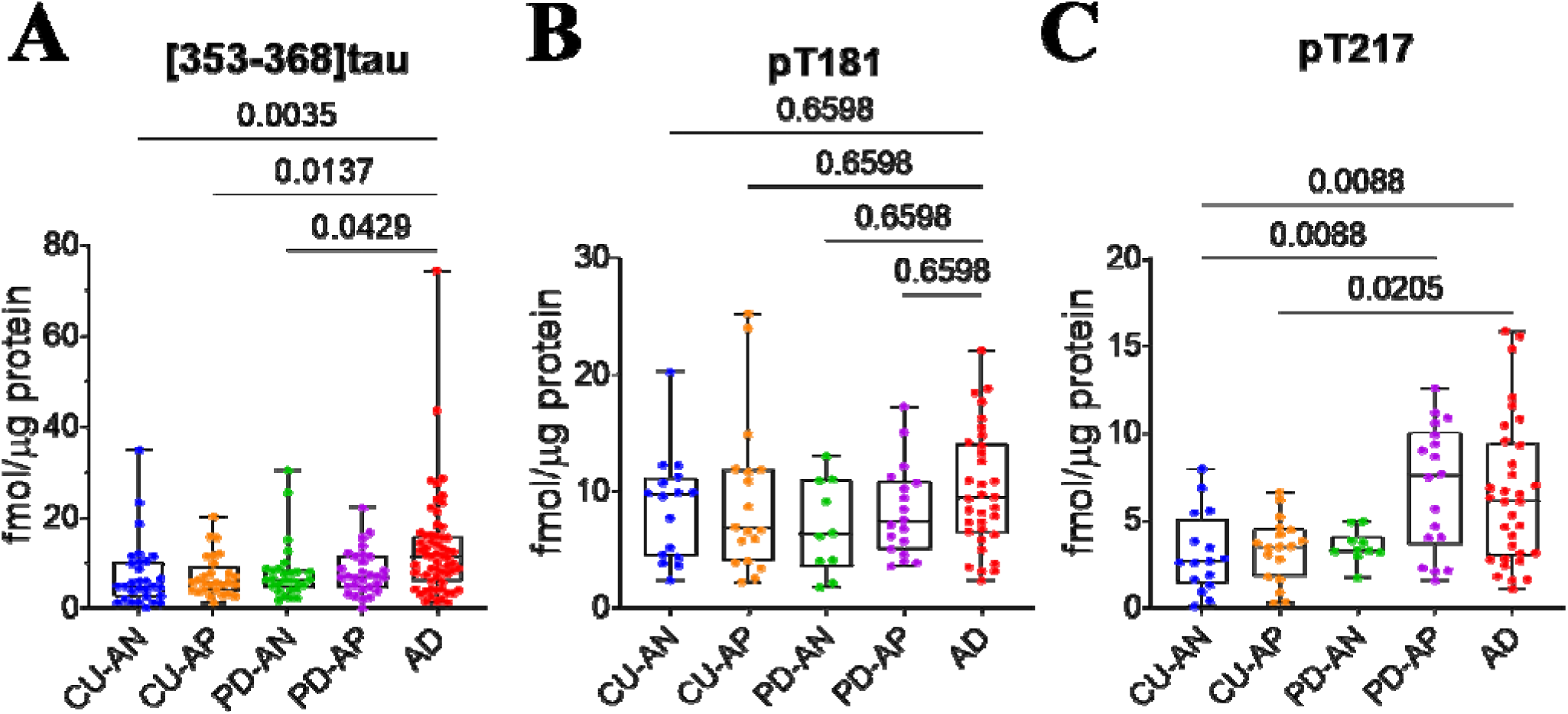
Box and whisker plots for the concentrations (fmol/µg protein) for MTBR tau species and phosphorylated tau (ptau) species (fmol/µg protein) in the detergent soluble brain homogenates. (A) MTBR R4 (residue 353-368) (B) pT181 and (C) pT217 species. Note MTBR tau and ptau pT217 was significantly increased in AD compared to CU-AP and CU-AN, while pT181 was unaffected. AD, Alzheimers disease (red); CU-AN, control brains without Aβ plaques (blue); CU-AP, control brains with Aβ plaques (orange); PD-AN, Parkinson’s disease (PD) brains without Aβ plaques (green); PD-AP, PD brains with Aβ plaques (purple). Statistically significant differences between the different groups are highlighted, Kruskal-Wallis test followed by Benjamini-Hochberg correction was performed for multiple comparisons.

### Isomerized A**β** ratios and their diagnostic performance

Aβ_1–15_ with two Asp residues (Asp-1 and Asp-7) can undergo individual isomerization/epimerization events, resulting in a total sixteen isomeric peptides from the combination of their L/D and/or iso-L/iso-D forms. We measured the isomerization of Asp-1 and Asp-7 on Aβ_1-15_ across the cohort by quantifying the four most abundant Aβ_1-15_ isomeric species found in AD.^17^ Previously, we have demonstrated major PTM on Aβ_1-15_ is isomerization of Asp-1 and Asp-7, with more than 85 % Aβ_1-15_ being isomerized compared to only 50 % isomerization in age matched control brains in amyloid-rich fractions of the cortical tissue.^17^ In this study, we quantified ratios of non-isomerized (native) Aβ_1-15_, and three isomerized species — mono-iso-Asp-L, di-iso-Asp-L and mono-iso-D-di-iso-Asp Aβ_1-15_ in the brains with fibrillar amyloid plaque pathology in the cohort. We observed native Aβ_1-15_ species (Asp1,7-L) in the AD brain (n=60) (Figure 6A) was significantly decreased (11.47 ± 7.21 %) compared to CU-AP (19.21 ± 9.78 %) (*p* = 0.0003) and PD-AP (20.65 ± 8.06 %) (*p* < 0.0001). We also observed mono-iso-Asp-L Aβ_1-15_ species was significantly decreased (Figure 6B) in AD brain (23.71 ± 4.49 %) compared to CU-AP (28.49 ± 6.18%) (*p* = 0.0007) and PD-AP (30.98 ± 5.69 %) (*p* < 0.0001). The doubly isomerized species, di-iso-Asp-L Aβ_1-15_ was significantly increased in AD brain (38.38 ± 6.70 %) compared to CU-AP (31.09 ± 10.59 %) (*p* = 0.0005) and PD-AP (30.60 ± 8.19 %) (*p* = 0.0002) (Figure 6C). In addition, we also observed the mono-iso-D-di-iso-Asp Aβ_1-15_ species was significantly increased in AD brain (11.51 ± 3.30 %) compared to CU-AP (7.17 ± 3.19 %) (*p* < 0.0001) and PD-AP (7.63 ± 2.52 %) (*p* < 0.0001) (Figure 6D). We observed no significant differences in the native as well as the isomerized species of Aβ_1-15_ between CU-AP and PD-AP.

**Figure 6:**
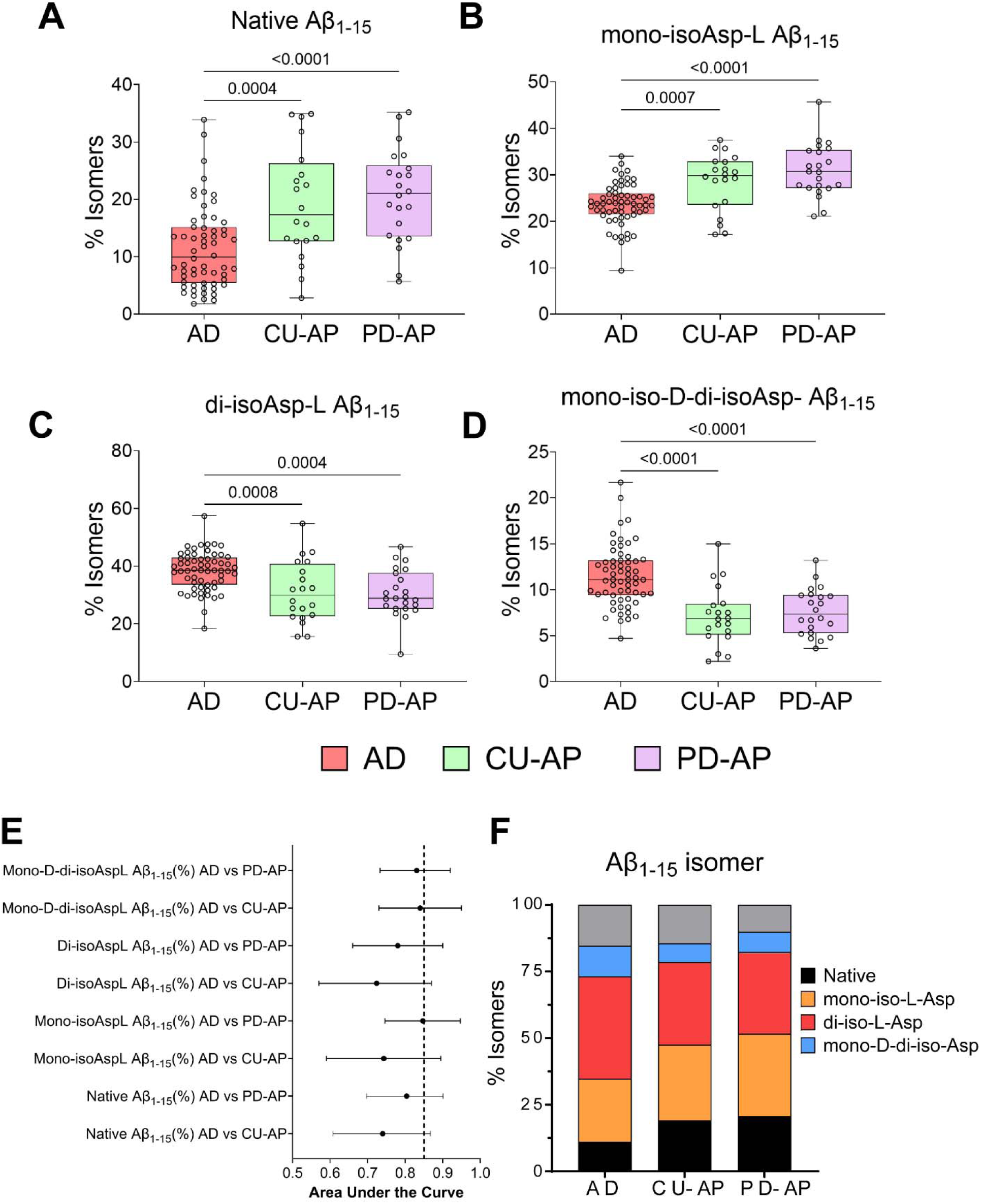
Isomerization of Aβ_1-15_ in detergent soluble fraction from human brain show distinct pattern across the AD neuropathological continuum. Box and whisker plots of percent isomers (%) of (A) native (unmodified Asp1,7-L) Aβ_1-15_ , (B) mono iso-Asp-L Aβ_1-15_, (C) doubly isomerized (di-iso-Asp-L) Aβ_1-15_ and (D) mono-iso-D-di-iso-Asp Aβ_1-15_ in AD, CU-AP and CU-AN. Statistically significant differences between the different groups are highlighted, Kruskal-Wallis test followed by Benjamini-Hochberg correction was performed for multiple comparisons. (E) Correspondence of isomer distribution in preclinical AD and non-AD dementia compared to AD. Points represent the ROC area under the curve (AUC) estimate (middle point) and the error bars represent the 95 % confidence intervals. Note the oldest doubly isomerized species distinguished AD from other comorbidities with high accuracy. (F) Bar graph of relative distribution of native (black), mono isomerized (orange),doubly isomerized (red) and mono-iso-D-doubly isomerized (blue) species of the total Aβ_1-15_ found in AD, CU-AP and PD-AP. Note while native (11 %) and mono-isomerized (24 %) Aβ_1-15_ contribution decreases in AD, the relative distribution of doubly isomerized (38 %) and mono-epimerized doubly isomerized (11 %) Aβ_1-15_ increases in AD compared other co-pathologies. Grey represents rest of the isomerized species of Aβ_1-15_. AD, Alzheimer’s disease; CU-AN, control tissue without Aβ plaques; CU-AP, non-demented control brain with Aβ plaques; PD-AN, PD brains without Aβ plaques; PD-AP, PD brains with Aβ plaques.

We further assessed the diagnostic performance of the isomers of Aβ_1-15_ in distinguishing AD from comorbidities with Aβ aggregates/deposits. We observed mono-iso-D-di-iso-Asp Aβ_1-15_ distinguished AD with high accuracy from CU-AP (AUC = 0.84 [0.73-0.95] and PD-AP (AUC = 0.83 [0.73-0.92] (Figure 6B, Figure S8). In addition, mono-iso-Asp-L Aβ_1-15_ distinguished AD with high accuracy from PD-AP (AUC = 0.85 [0.75-0.95] and moderately high accuracy from CU-AP (AUC = 0.74 [0.59-0.89]. The doubly isomerized Aβ_1-15_ species distinguished AD with high accuracy from PD-AP (AUC = 0.78 [0.66-0.90] and moderately high accuracy from CU-AP (AUC = 0.72 [0.57-0.87]. Native Aβ_1-15_ species also distinguished AD from PD-AP (AUC = 0.80 [0.69-0.90] and CU-AP (AUC = 0.74 [0.61-0.87].

Finally, we compared the fractional isomerization of Aβ_1-15_ with the AD neuropathologic changes (Figure 7), and observed the native (non-isomerized) Aβ_1-15_ was negatively correlated with the Thal amyloid phase (r = -0.46, *p* < 0.001, A score), Braak tau stages (r = -0.51, *p* < 0.001, B score) and neuritic plaque scores (r = -0.46, *p* < 0.001, C score) (Figure 7A-C and Figure S8). The mono-isomerized (Aβ_1-15_ with either Asp-1 or Asp-7 residues are isomerized) Aβ_1-15_ was negatively correlated with the Thal amyloid phase (r = - 0.51, *p* < 0.001, A score), Braak tau stages (r = -0.57, *p* < 0.001, B score) and neuritic plaque scores (r = -0.51, p < 0.001, C score) (Figure 7D-F). Whereas the doubly isomerized Aβ_1-15_ species (di-iso-Asp-L Aβ_1-15_) was positively correlated with Thal amyloid phase (r = 0.43, *p* < 0.001, A score), Braak tau stages (r = 0.48, *p* < 0.001, B score) and neuritic plaque scores (r = 0.43, *p* < 0.001, C score). The oldest doubly isomerized Aβ_1-15_ species (mono-iso-D-di-iso-Asp Aβ_1-15_) showed the best positive correlation with Thal amyloid phase (r = 0.54, *p* < 0.001, A score), Braak tau stages (r = 0.56, *p* < 0.001, B score) and neuritic plaque scores (r = 0.55, *p* < 0.001, C score) (Figure 7G-K and Figure S8). Braak tau stages and neuritic plaque scores. These results indicate the native/unmodified (Asp-1 and Asp-7) Aβ_1-15_ and mono-isomerized Aβ_1-15_ are more abundant in intermediate ADNC, decreasing significantly in late ADNC, while more isomerized/older Aβ_1-15_ species becomes abundant in late ADNC stages (Figure 7).

**Figure 7.**
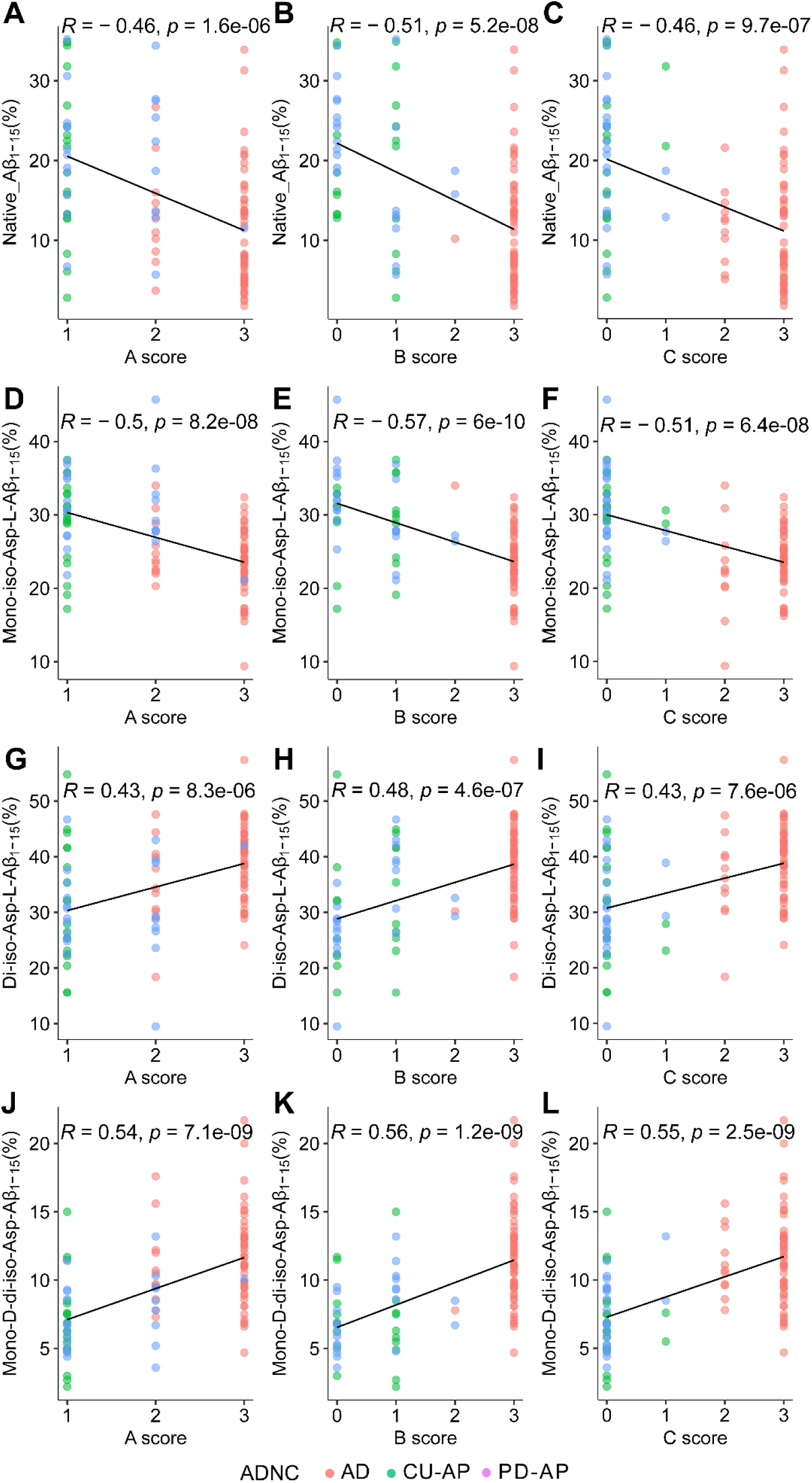
Pearson correlation analyses between the isomerized species (%) of Aβ_1-15_ with neuropathological AD changes. (A-C) Native Aβ_1-15_, (D-F) mono isomerized Aβ_1-15_, (G-I) doubly isomerized Aβ_1-15_ (di-iso-Asp-L) and (J-L) mono-iso-D-di-iso-Asp compared with Thal amyloid phases (A scores), Braak tau stages (B scores) and CERAD scores (C scores). Moderate negative correlation was observed for the native Aβ_1-15_ and mono-isomerized Aβ_1-15_ with AD neuropathologic changes (ADNC), while positive strong correlations were observed for doubly isomerized (di-iso-Asp and mono-D-di-iso-Asp) Aβ_1-15_ species with the ADNC. AD, Alzheimer’s disease; CU-AN, control brains without Aβ plaques; CU-AP, control brains with Aβ plaques; PD-AN, PD brains without Aβ plaques; PD-AP, PD brains with Aβ plaques.

## Discussion

In this study, we present the quantitative analysis of multiple PTMs associated with Aβ in human brain tissue in AD and non-AD dementia that have amyloid plaques as comorbidities and compared our measures with their postmortem neuropathological scores. Our study used a large human brain sample set (n=176) in combination with targeted mass spectrometry without any affinity enrichment (immunoprecipitation). This provided an unbiased representation of detergent soluble fibrillar Aβ aggregates and the most prominent PTMs, without specifically enriching any conformations/species of Aβ. The major findings of our study are (i) isomerization on Asp-1 and Asp-7 is the most common PTM on fibrillar Aβ aggregates found in symptomatic AD, preclinical AD and non-AD dementia, (ii) percent isomerization (isomer ratio) of Asp-1 and Asp-7 on fibrillar Aβ aggregates, extractable from cortical tissue, can distinguish between intermediate ADNC and late ADNC stages with high accuracy and, and (iii) citrullinated Aβ_3pGlu_ was associated with late ADNC, being significantly elevated in AD compared to non-AD dementia or preclinical AD cases with Aβ plaque pathology.

Long-lived proteins/peptides are subject to spontaneous chemical modifications and iso-aspartate formation is one of the most common PTM detected in long-lived proteins.^37,38^ Iso-Asp Aβ modification was identified by Roher more than 30 years ago from AD brain parenchymal plaques.^24^ Structural reorganization due to Asp isomerization could make the fibrillar Aβ inherently resistant to enzymatic degradation by the primary cathepsins found in the lysosomes.^39^ We have previously estimated the age of isomerized Aβ present in human brains using the *in vitro* half-life of Aβ Asp isomerization (*t*_1/2_ 231 days, Asp-1; *t*_1/2_ 432 days, Asp-7).^17,39^ AD brains with almost 90 % isomerized Aβ_1-15_ indicated the age of this peptide is more than 4 years, in-line with our previous estimates. Whereas the non-demented controls positive for Aβ plaques with 80 % isomerized Aβ_1-15_ indicate the Aβ aggregates in these brains are 3 years old. This suggests the age of the Aβ plaques in AD is more advanced/older compared to non-AD comorbidities. It is worth noting these estimates don’t include the brain Aβ biogenesis and altered clearance rates observed in cerebral amyloidosis participants,^13,14^ that would push the age estimates of Aβ aggregates in the plaques into decades. Interestingly, racemization rates of Asp-L to Asp-D used by Muller-Hill and Beyreuther to study amyloid plaque cores estimated the age of plaques at 30 years.^40^ *In vitro* estimation of the exact isomerization rates along with the modelling of the clearance rates will provide better estimates of the age of amyloid plaques in AD compared to preclinical AD.

Recent study using both immunohistochemistry and biochemical analysis found abundant isoAsp-7 Aβ in AD followed by other dementias such as Lewy body dementia, vascular dementia and preclinical AD.^27^ Our results indicate that not all isomerized Aβ species are increased in AD, relative contribution of the unmodified (native) and early mono-isomerized Aβ (either isoAsp-1 or isoAsp-7) actually decreases in symptomatic AD compared to preclinical AD and non-AD dementia. While the doubly isomerized (isoAsp1 and isoAsp-7) increase in older/mature parenchymal plaques in symptomatic AD. The conversion of early mono-isomerized species to older doubly-isomerized/epimerized Aβ also correlated with the neuropathological staging (ABC scores). Does this mean fibrillar Aβ once aggregated/deposited into the extracellular plaques remain quiescent and are subject to spontaneous chemical modifications without any turnover — a question that will need further investigation using postmortem human brain tissue samples from participants that had undergone stable isotope labelling kinetics (SILK) during life.^41,42^ IsoAsp-7 Aβ when administered peripherally to transgenic mice model induced cerebral amyloid pathology and have been found to be neurotoxic to cell cultures compared to native Aβ.^43,44^ Iso-aspartate accumulation is known to be lethal in the PIMT (protein iso-aspartate methyltransferase) deficient mouse, suffering from epileptic seizures.^45^ PIMT has shown neuroprotective role by reducing the hydrophobicity and toxicity of the Aβ oligomers.^46^ Interestingly, targeting isomerized Aβ using a monoclonal antibody targeting isoAsp-7 demonstrated reduction in amyloid pathology and ameliorated behavioral deficits in 5xFAD mice.^47^ These results support specifically targeting the older isomerized Aβ for clearance will be better strategy for immunotherapy.

N-terminus of the Aβ extracted from parenchymal plaques is known to demonstrate high heterogeneity.^48–50^ Such heterogeneity has been proposed to be derived from the activities of multiple proteases on the canonical Aβ_42_ produced from APP.^4,51,52^ Another plausible mechanism is spontaneous non-enzymatic process, that leads to sequential truncations (“laddering”) of the Aβ terminus.^53^ This is supported by the flexibility of the N-terminus from Cryo-EM data of human brain derived fibrillar Aβ, the ordered core of each protofilament includes Aβ residues from Gly9 to Ala42, with the N-terminus forming the fuzzy coat of the Aβ filaments.^30^ Historically, the first sequenced parenchymal Aβ described by Masters was the truncated species Aβ_4-42_.^1^ The presence of this peptide have been demonstrated by us and others using mass spectrometry in AD brain in both the parenchyma as wells as vascular deposits in familial mutation carriers.^17,50,54,55^ The origin of the Aβ(4-x) peptide in the plaques have been a mystery and only recently ADMATS4, an extracellular metalloprotease, has been shown to enzymatically cleave and generate Aβ with Phe4 at the N-terminus.^56^ Pyroglutamate modified Aβ (Aβ_3pGlu_) is another common modified Aβ that was first described by Saido and Roher, and has been demonstrated to correlate with AD neuropathologic changes.^57,58^ Aβ_3pGlu_ has emerged as promising pharmacologic target in AD therapy, with the recent U.S. Food and Drug administration (FDA) approval of Donanemab for the treatment for early symptomatic AD.^59^ Both of these peptides have increased propensity to aggregate and high neurotoxicity, consequence of change on the biophysical properties due to the loss of charged residues at the N-terminus.^60^ Our data demonstrate that much of the N-terminus has Asp1 (∼ 60%), while Aβ with pGlu3 and Phe4 as N-termini that can be extracted from the parenchymal Aβ plaque accounts for 15 % and 20 %, respectively. These relative distributions are not significantly altered in AD compared to control brains with Aβ amyloid pathology (Figure 3), indicating these modified Aβ are normal constituents of parenchymal Aβ plaques. Most importantly, irrespective of the origin of these truncations, these truncated peptides are abundant in early/immature plaques in preclinical AD and non-AD dementia and targeting them for removal from the brain early in course of the disease would be therapeutically promising.

We recently described a novel PTM on Aβ, deimination of the Arg5 residue — citrullination and found pyroglutamate Aβ is hypercitrullinated in AD.^21^ Our results suggest that citrullinated Aβ_3pGlu_ is significantly increased only in AD and can discriminate preclinical AD and non-AD dementia. Citrullination is well known feature of inflammation and has been linked to increased neuroinflammation in AD.^61,62^ Altered peptidyl arginine deiminase (PAD) activity has been linked to AD.^63^ Our data highlights that increased citrullinated Aβ_3pGlu_ is associated with fibrillar Aβ plaques in late symptomatic AD, not evident in earlier/diffuse plaques in preclinical AD and non-AD dementia. Another strength of our study is the evaluation of phospho tau (pT217) and total tau (MTBR-tau) levels in a subset of this cohort. Recent biomarker developments have highlighted importance of these brain derived tau species in the biofluids (CSF/blood) in identifying and monitoring neurodegenerative progression in AD.^64–67^ Our findings confirm that phospho tau levels are already elevated in intermediate ADNC stages in the cortical region, alongside with phospho tau co-pathology in dementia with Lewy bodies (DLB).^68,69^ However, detergent soluble pT217 levels showed modest diagnostic performance in separating late AD stages compared to co-pathologies. Based on the diagnostic performances of the brain Aβ PTMs including Aβ_3pGlu_, citrullinated Aβ_3pGlu_, and Asp-1 and Asp-7 isomerization on Aβ may provide an opportunity to distinguish comorbidities and disease progression if detected in biofluids such as cerebrospinal fluid (CSF) or blood in preclinical AD or mid-cognitively impaired AD patients. Along with targeting Aβ_3pGlu_, therapeutic targeting of citrullinated Aβ_3pGlu_ and inhibition of PAD activity could be promising in treating late stages of the disease.

In this study, we also quantitatively estimated the Aβ_42_ and Aβ_40_ in the detergent soluble brain fraction. We choose to investigate urea-detergent fraction in this study as we have previously demonstrated that most of the brain Aβ is still partitioned in the membranous (57 %) and insoluble fully polymerized phase fraction (∼ 41 %).^17,70^ In line with what we and others have previously found,^15,17,33,57,70–72^ Aβ_42_ is the most abundant species (∼ 90 %) of the total fibrillar Aβ aggregates depositing in the parenchyma. We observed, Aβ_40_ was significantly increased in AD and distinguished late ADNC with high accuracy compared to intermediate ADNC. This has been previously reported using both immunohistochemistry as well as MALDI-MS imaging, where mature/cored plaques recruit Aβ_40_ over time.^16,55,73^ While Aβ_42_ is the main Aβ species in the parenchymal plaques, comparatively less hydrophobic Aβ_40_ is the predominant species in the vascular amyloidosis observed in cerebral amyloid angiopathy (CAA). ^74–76^ The exact mechanism of the preferential deposition of two different Aβ isoforms in these distinct compartments of the brain needs further elucidation. Diffusion of Aβ_40_ across the perivascular drainage pathways leading to its deposition in the vessel walls compared to deposition of more hydrophobic Aβ_42_ in the brain parenchyma has been postulated as a possible molecular mechanism.^71,77^ Hence anti-amyloid therapies that target the C-terminus of the Aβ, specifically targeting Aβ_42_ for clearance could better maneuver the amyloid equilibrium in the AD brain. This is more crucial, as lower rates of anti-amyloid related imaging abnormalities (ARIA) have been found in clinical trials that used antibodies that target mid-peptide and C-terminal regions of Aβ,^78,79^ unlike trials that target N-terminal or conformational forms of Aβ for clearing plaques.^31,32,80^ Targeting the most abundant and old isomerized Aβ for effector mediated elimination would reduce the therapeutic drug amount along with treatment related adverse side effects.

## Conclusion

Post translationally modified Aβ play a crucial role in aggregation, inhibiting degradation by lysosomal enzymes that results in accumulation of fibrillar assemblies that are highly neurotoxic. We hypothesized that distinct patterns of modified Aβ variants could distinguish late-stage neuropathologic changes in AD from early deposits observed in preclinical AD. Indeed, we report that the mono isomerized Aβ are abundant species in preclinical-AD and non-AD dementia with Aβ plaque pathology, while older doubly isomerized Aβ species are increased in AD. We also discovered that citrullinated Aβ_3Glu_ distinguished AD from non-AD dementia. Our results indicate that targeting older isomerized Aβ in AD would be better pharmacological approach in AD treatment. Furthermore, our mass spectrometry platform offers an invaluable tool for investigating changes in pathological burden in postmortem tissues, providing insights into how anti-amyloid targeting therapeutics impact disease progression beyond alterations in imaging and biofluid biomarkers.

## Authors’ contributions

S.M., C.L.M, and B.R.R. conceived the study. S.M. designed and performed the proteomics experiments, tandem mass spectrometry statistical analysis and interpretation of the data. C.M. and F.H. provided brain bank access, neuropathological assessments and diagnostics. S.M., C.D. and K.P. performed brain homogenization and urea detergent extraction. The first draft of the manuscript was written by S.M., C.L.M. and B.R.R. The manuscript was written with input from all the authors. All authors have read and approved the manuscript.

## Acknowledgements

We thank Geoffrey Pavey from the Victorian Brain Bank, Ian Birchall, Fairlie Hinton, Liang Jin, Stephan Klatt and Kevin J. Barham from the Florey Institute for valuable discussions and help during various stages of analysis. We would like to thank the Florey Institute Neuroproteomics Facility, and the Melbourne Mass Spectrometry and Proteomics Facility (MSPF). S.M. and R.C. would also like to thank the support the resources and effort provided by the Tracy Family Stable Isotope Labeling Quantitation Center at the Department of Neurology, Washington University in St. Louis.

## Data availability statement

The quantitative data are available as supplementary materials accompanying this article. Additional data and materials supporting the findings of the study can be requested from the corresponding authors.

## Authors’ information

Soumya Mukherjee

Celine Dubois

Keyla Perez

Catriona McLean

Colin L. Masters

Blaine R. Roberts

