## Supplementary Figures for "Isomerized Aβ in the brain can distinguish the status of amyloidosis in the Alzheimer’s Disease spectrum"

Soumya Mukherjee^1,2^, Reid Coyle,^1,2^ Celine Dubois^3^, Keyla Perez^3^, Ian Birchall,^3^ Fairlie Hinton,^3^ Catriona McLean,^3,4^ Colin L Masters^3^, Blaine R Roberts^5,6^

^1^The Tracy Family SILQ Center, Washington University School of Medicine in St. Louis, MO, USA

^2^Department of Neurology, Washington University School of Medicine in St. Louis, MO, USA

^3^The Florey Institute of Neuroscience and Mental Health, Parkville, Victoria, Australia

^4^Department of Anatomical Pathology, Alfred Hospital, Prahran, Victoria 3004, Australia

^5^Department of Biochemistry, Emory University, GA, USA

^6^Department of Neurology, Emory University, GA, USA

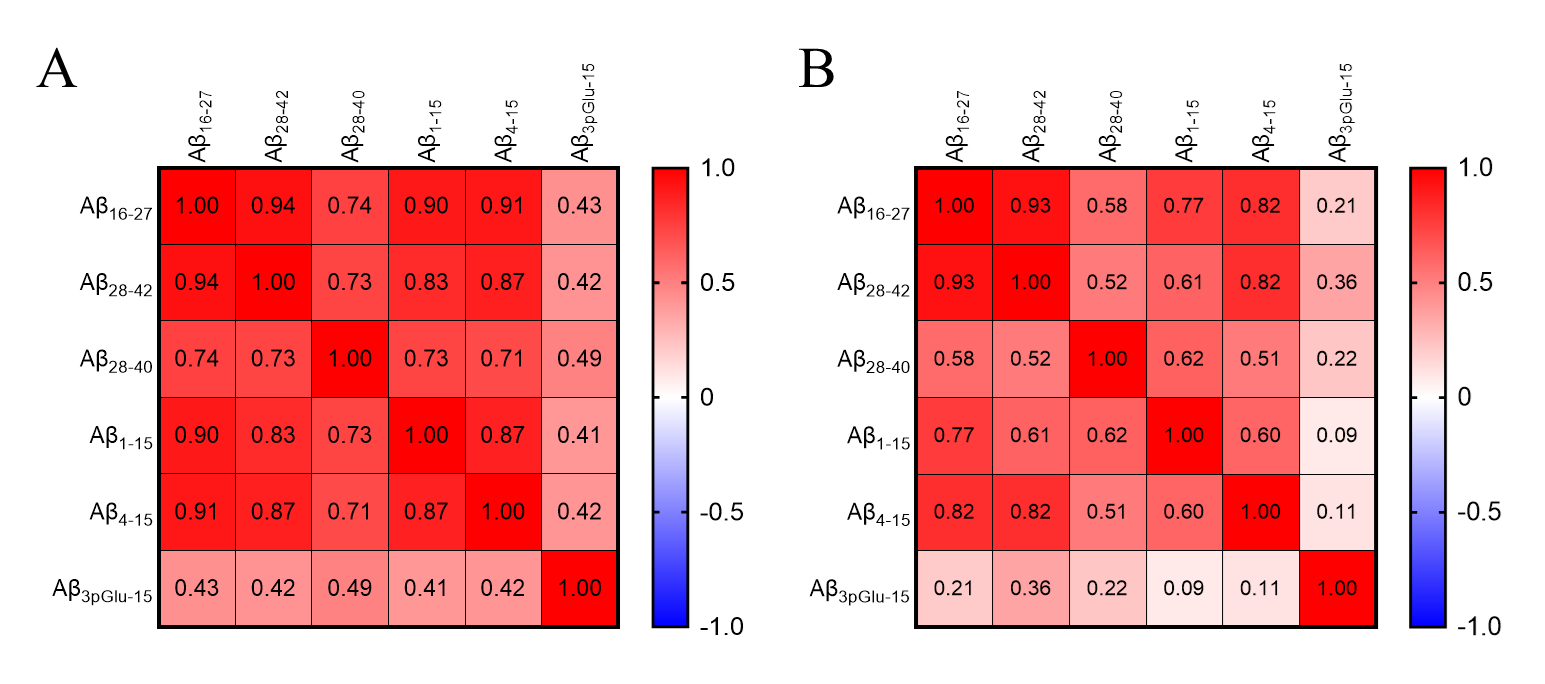

**Supporting Figure S1**. Correlation matrix presenting Spearman’s correlations for the Aβ peptide concentrations across the (A) whole cohort and (B) AD brain homogenates. High correlation between Aβ_42_ and Aβ_16-27_ is found (r > 0.9), while Aβ_40_ has relatively lower correlation with both the Aβ_42_ (r ~ 0.5) and Aβ_16-27_ (r ~ 0.6) in AD brains. N-truncated Aβ_4-15_ has higher correlation with the Aβ_42_ and Aβ_16-27_ (r = 0.82), compared to Aβ_1-15_ that has lower correlations with the Aβ_42_ (r ~ 0.8) and Aβ_16-27_ (r ~ 0.6) in AD brains. Aβ_3pGlu-15_ has relatively low correlation with N-terminus Aβ_1-15_ (r ~ 0.1) and Aβ_4-15_ (r ~ 0.11).

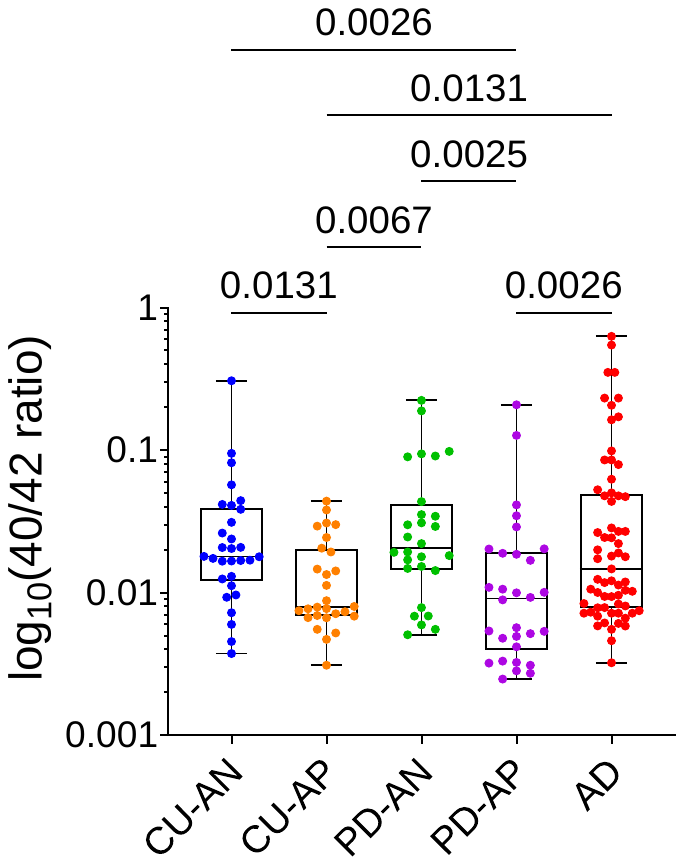

**Supporting Figure S2.** Box and whiskers plot of log transformed Aβ_40/42_ ratio in the detergent soluble brain fractions. All the individual brain samples are depicted as filled circles as the following: blue, CU-AN; orange, CU-AP; green, PD-AN; violet, PD-AP and red, AD. Statistical significance was determined by two-way ANOVA with multiple testing comparisons. Abbreviations: AD, Alzheimer’s disease; CU-AN, non-demented control tissue without Aβ plaques; CU-AP, non-demented control brain with Aβ plaques; PD-AN, PD brains without Aβ plaques; PD-AP, PD brains with Aβ plaques.

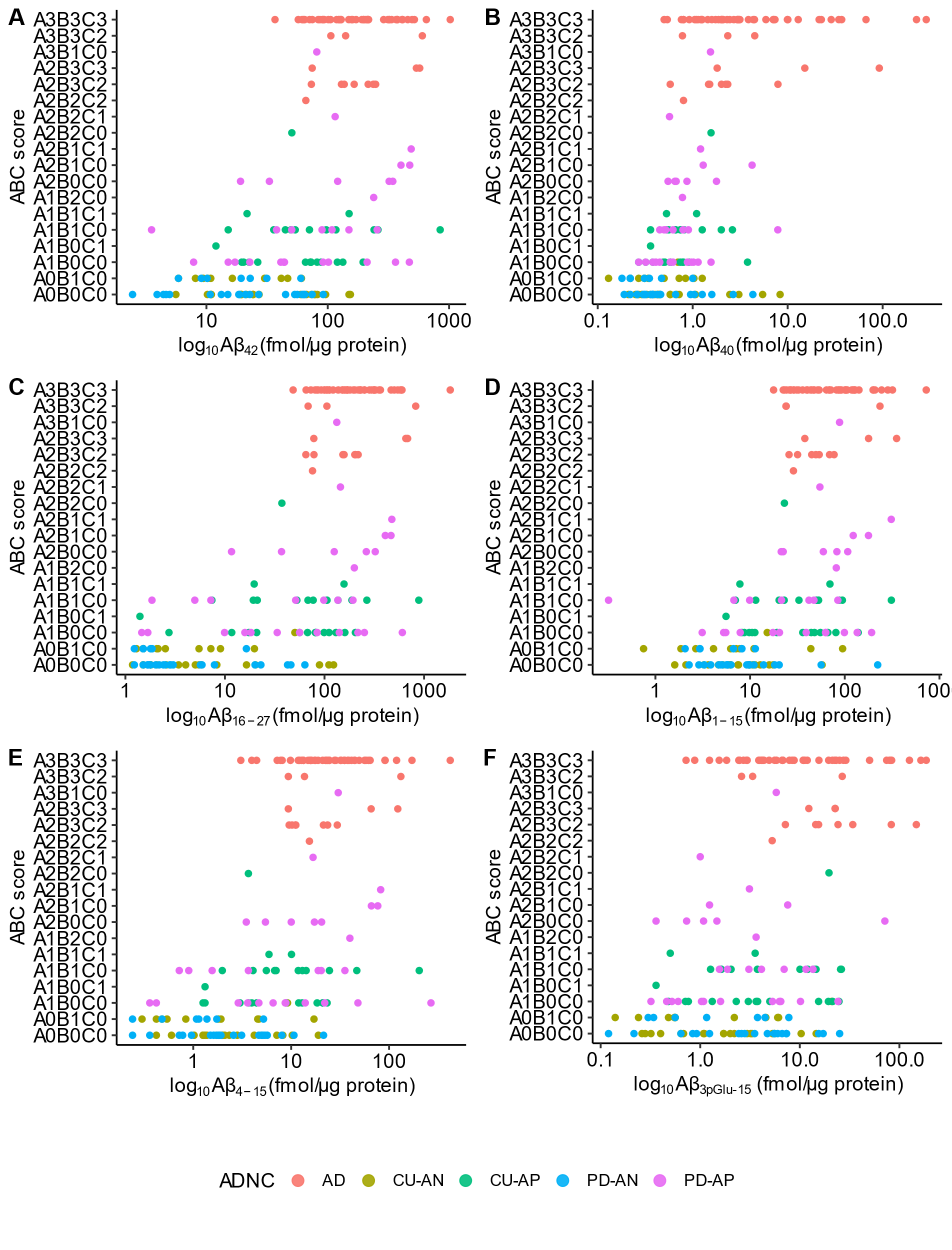

**Supporting Figure S3.** Cross-sectional associations between Aβ isoforms in the brains and the AD neuropathological change. Log transformed concentrations (fmol/µg protein) of (A) Aβ_42_, (B) Aβ_40_, (C) Aβ_16-27_, (D) Aβ_1-15_, (E) Aβ_4-15_ and (F) Aβ_3pGlu-15_ are compared with the ABC scores. Abbreviations: AD, Alzheimer’s disease; CU-AN, non-demented control tissue without Aβ plaques; CU-AP, non-demented control brain with Aβ plaques; PD-AN, PD brains without Aβ plaques; PD-AP, PD brains with Aβ plaques.

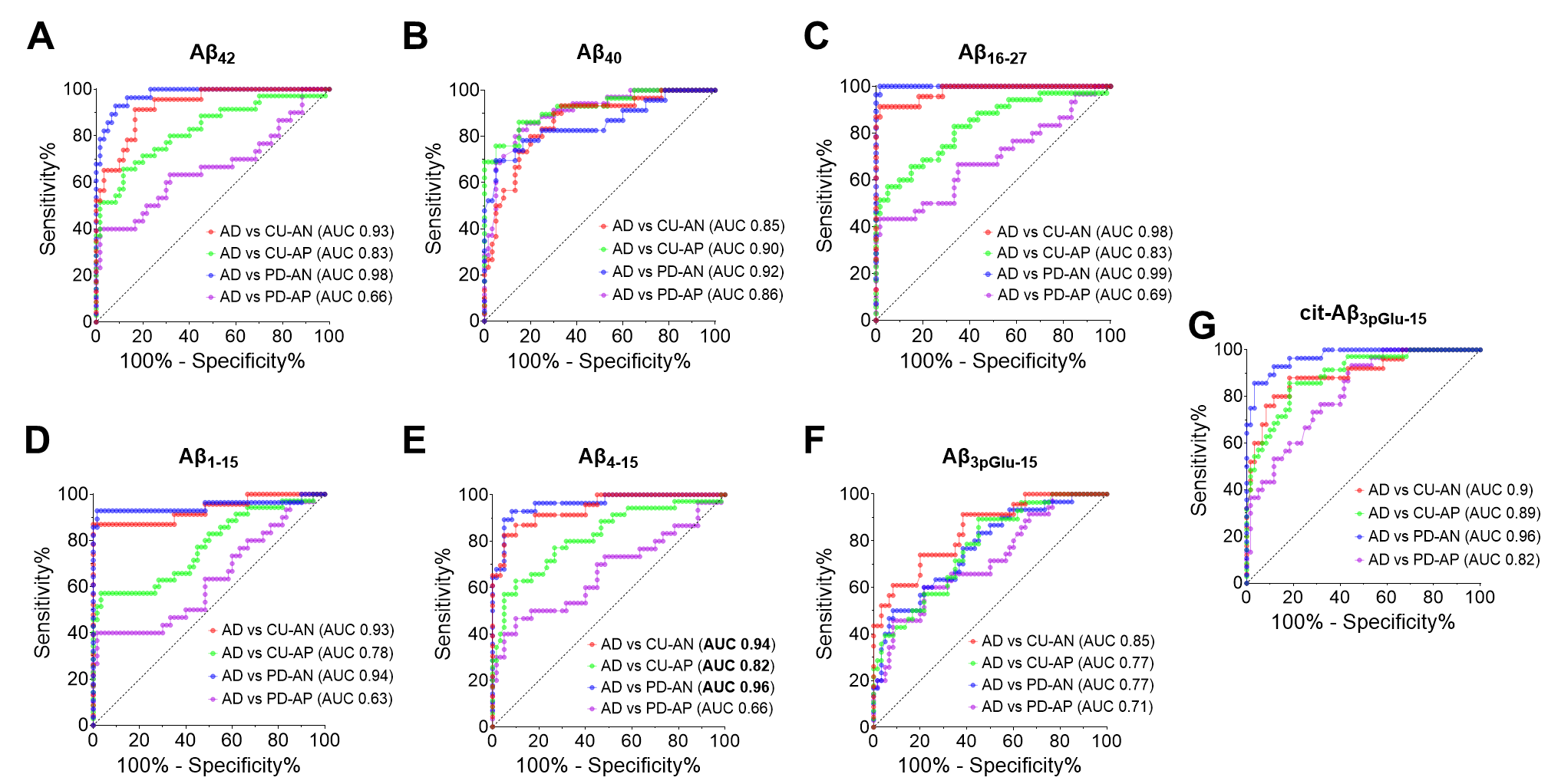

**Supporting Figure S4**. Receiver operating characteristic (ROC) curves for (A) Aβ_42_, (B) Aβ_40_, (C) Aβ_16-27_, (D) Aβ_1-15_, (E) Aβ_4-15_, (F) Aβ_3pGlu-15_ and (G) cit-Aβ_3pGlu-15_ concentrations (fmol/µg protein) in the detergent soluble brain homogenates with their respective area under the curve (AUCs) in differentiating AD from CU-AN (red), CU-AP (green), PD-AN (blue) and PD-AP (purple). Abbreviations: AD, Alzheimer’s disease; CU-AN, non-demented control tissue without Aβ plaques; CU-AP, non-demented control brain with Aβ plaques; PD-AN, PD brains without Aβ plaques; PD-AP, PD brains with Aβ plaques.

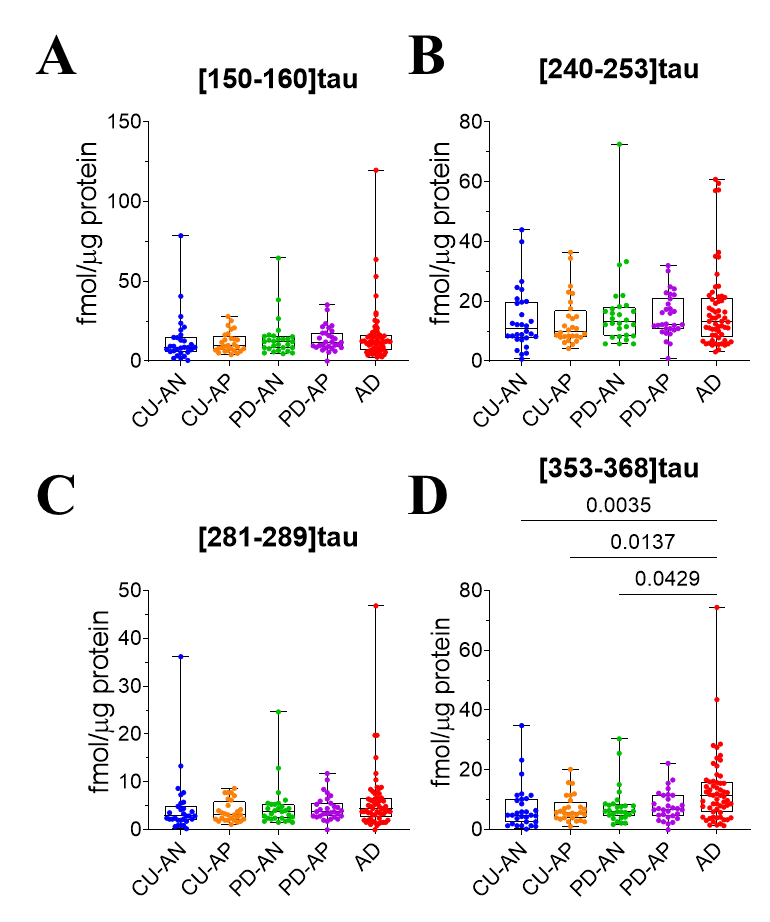

**Supporting Figure S5.** Detergent soluble MTBR-tau is increased in late ADNC. Box and whisker plots for (A) PRR tau (residue 150-160), (B) MTBR240-tau (residue 240-253, R1), (C) MTBR281-tau (residue 281-289, R2) and (D) MTBR353-tau (residue 353-368, R4) concentrations (fmol/µg protein) in the detergent soluble brain homogenates. Abbreviations: AD, Alzheimer’s disease; CU-AN, non-demented control tissue without Aβ plaques; CU-AP, non-demented control brain with Aβ plaques; PD-AN, PD brains without Aβ plaques; PD-AP, PD brains with Aβ plaques.

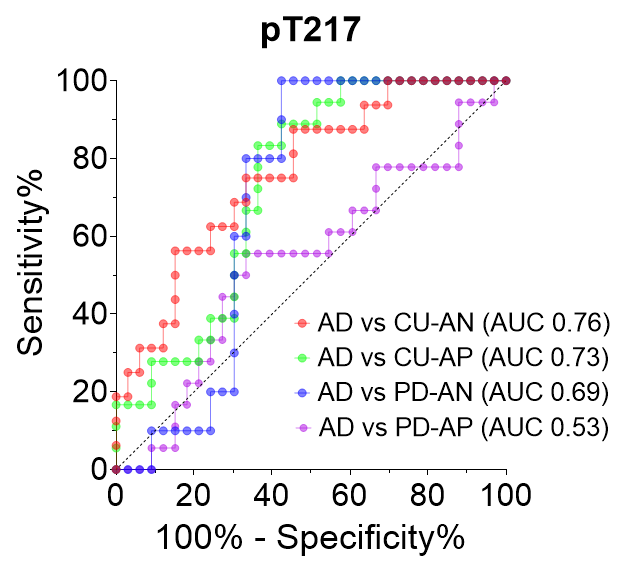

**Supporting Figure S6**. Phosphorylated tau-T217 is increased in both intermediate and late ADNC stages. Diagnostic specificity and sensitivity between AD and other comorbidities from the ROC curves for pT217. The AUCs are highlighted, which indicate diagnostic performance of pT217 is low between preclinical AD, PD-AP and symptomatic AD. Abbreviations: AD, Alzheimer’s disease; CU-AN, non-demented control tissue without Aβ plaques; CU-AP, non-demented control brain with Aβ plaques; PD-AN, PD brains without Aβ plaques; PD-AP, PD brains with Aβ plaques.

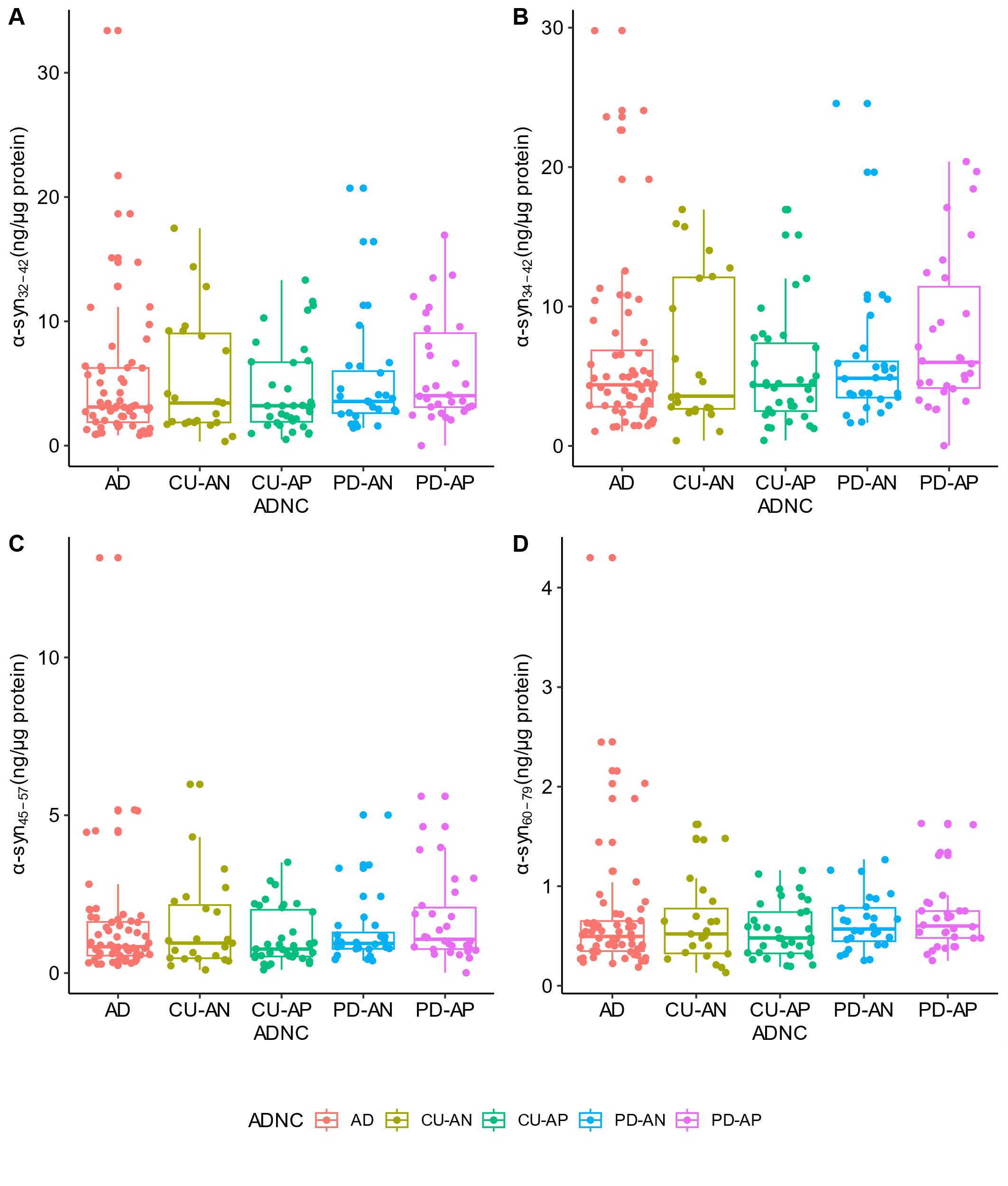

**Supporting Figure S7.** Detergent soluble α-syn concentrations does not change across different ADNC. Box and whisker plot of (A) α-syn_32-42_, (B) α-syn_34-42_, (C) α-syn_45-57_ and (D) α-syn_60-79_ concentrations (ng/µg protein) in the detergent soluble brain homogenates. Abbreviations: AD, Alzheimer’s disease; CU-AN, non-demented control tissue without Aβ plaques; CU-AP, non-demented control brain with Aβ plaques; PD-AN, PD brains without Aβ plaques; PD-AP, PD brains with Aβ plaques.

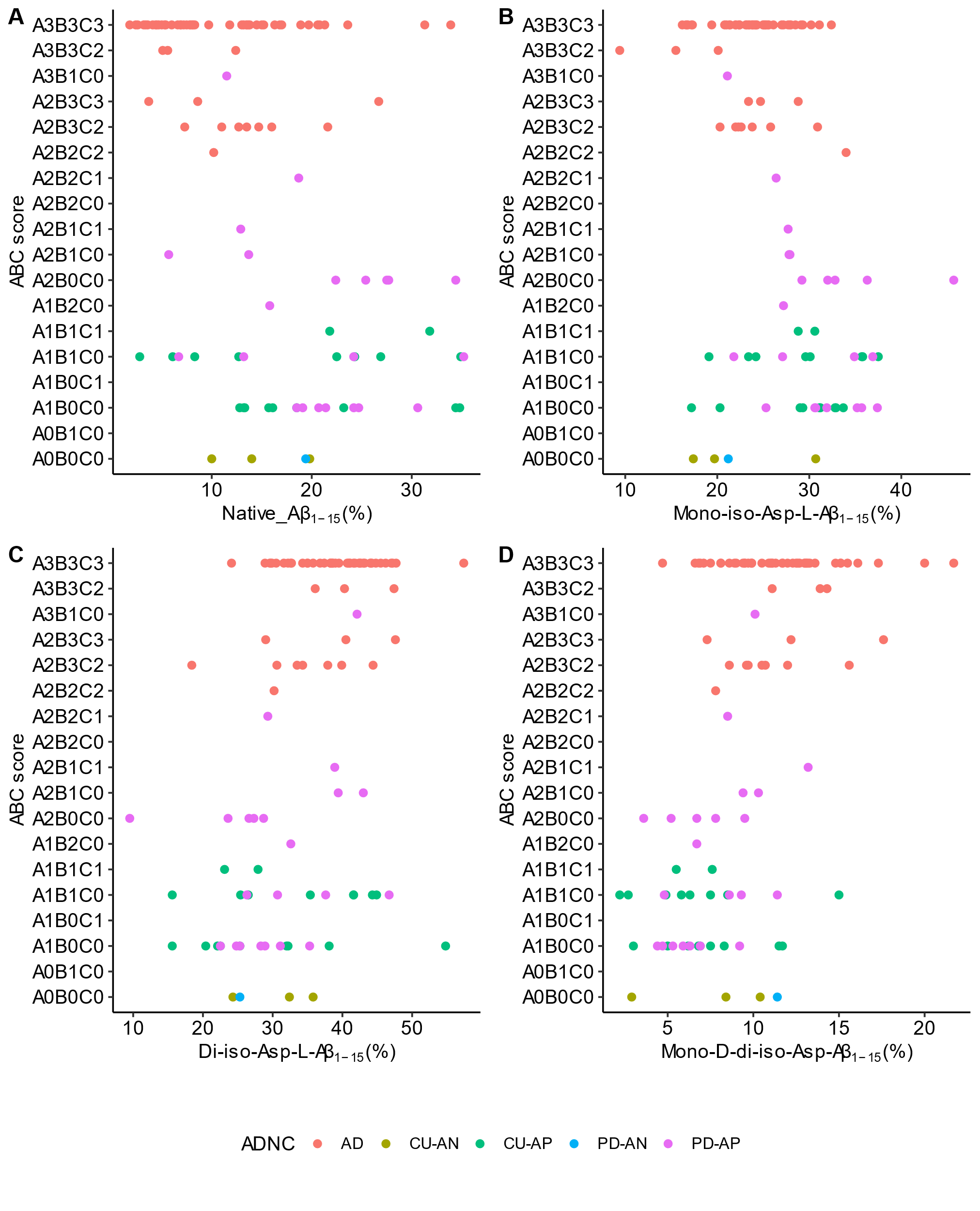

**Supporting Figure S8**. Cross-sectional associations between isomerized Aβ_1-15_ species in the brains and the AD neuropathological change. Percent ratios of (A) native Aβ_1-15_, (B) mono-iso-Asp Aβ_1-15_, (C) di-iso-Asp Aβ_1-15_ and (D) mono-D-di-iso-Asp Aβ_1-15_ ratios (%) are compared with the ABC scores of individual participants. Abbreviations: AD, Alzheimer’s disease; CU-AN, non-demented control tissue without Aβ plaques; CU-AP, non-demented control brain with Aβ plaques; PD-AN, PD brains without Aβ plaques; PD-AP, PD brains with Aβ plaques.
